# Potentiated cholinergic and corticofugal inputs support reorganized sensory processing in the basolateral amygdala during auditory threat acquisition and retrieval

**DOI:** 10.1101/2023.01.31.526307

**Authors:** Meenakshi M. Asokan, Yurika Watanabe, Eyal Y. Kimchi, Daniel B. Polley

**Affiliations:** Eaton-Peabody Laboratories, Massachusetts Eye and Ear, Boston MA 02114 USA; Division of Medical Sciences, Harvard Medical School, Boston MA 02114 USA; Department of Otolaryngology - Head and Neck Surgery, Harvard Medical School, Boston MA 02114 USA; Department of Neurology, Massachusetts General Hospital, Boston, MA 02114, USA

**Keywords:** temporal association area, higher-order auditory cortex, basolateral amygdala complex, corticoamygdalar neurons, corticofugal projections, discriminative fear learning, Pavlovian conditioning, pupil, facial motion, spike-triggered local field potential, acetylcholine, cholinergic modulation

## Abstract

Reappraising neutral stimuli as environmental threats reflects rapid and discriminative changes in sensory processing within the basolateral amygdala (BLA). To understand how BLA inputs are also reorganized during discriminative threat learning, we performed multi-regional measurements of acetylcholine (ACh) release, single unit spiking, and functional coupling in the mouse BLA and higher-order auditory cortex (HO-AC). During threat memory recall, sounds paired with shock (CS+) elicited relatively higher firing rates in BLA units and optogenetically targeted corticoamygdalar (CAmy) units, though not in neighboring HO-AC units. Functional coupling was potentiated for descending CAmy projections prior to and during CS+ threat memory recall but ascending amygdalocortical coupling was unchanged. During threat acquisition, sound-evoked ACh release was selectively enhanced for the CS+ in BLA but not HO-AC. These findings suggest that phasic cholinergic inputs facilitate discriminative plasticity in the BLA during threat acquisition that is subsequently reinforced through potentiated auditory corticofugal inputs during memory recall.

## Introduction

Thriving, if not merely surviving, requires a well-calibrated risk management system to evaluate potential threats in the environment and deploy adaptive behavioral responses. Threat evaluation has been modeled with a Pavlovian auditory fear conditioning paradigm in which a neutral sound is subsequently paired with an aversive stimulus, producing defensive behaviors (e.g., freezing) and heightened autonomic responses (e.g., pupil dilation) elicited by the auditory conditioned stimulus (CS). Associative memory of the threatening is sound is encoded by reorganized CS processing, synaptic plasticity, and genetic modifications in a distributed network of brain regions, though the basolateral amygdala complex (BLA) is widely understood to be an essential hub in this network (Herry and Johansen, 2014; Janak and Tye, 2015; LeDoux, 2007; Maren and Quirk, 2004; Tovote et al., 2015).

Auditory information reaches the BLA complex (identified here as the lateral, basal and basomedial amygdala) via descending neocortical projections as well as thalamic projections from the intralaminar nucleus and medial subdivision of the medial geniculate body (Barsy et al., 2020; Dalmay et al., 2019; Ledoux,’ et al., 1990; Romanski and Ledoux, 1993). The BLA is also densely innervated by cholinergic afferents from the basal forebrain (Gielow and Zaborszky, 2017; Mesulam et al., 1983; Woolf and Butcher, 1982). Although associative strengthening of the auditory CS response is observed in the thalamic (Belén Pardi et al., 2020; Edeline and Weinberger, 1992; Taylor et al., 2021), cortical (Weinberger, 2004), and cholinergic inputs to the BLA (Jiang et al., 2016; Likhtik and Johansen, 2019), these regions also receive feedback projections from the BLA (Aizenberg et al., 2019; Chavez and Zaborszky, 2016; Yang et al., 2016), thereby making the necessary involvement of associative CS plasticity in these regions uncertain, at least for the case where relatively simple auditory stimuli are used as a CS (e.g., a tone burst). Discriminative threat conditioning (DTC) can be distinguished from the broader class of Pavlovian auditory fear conditioning protocols by the use of repeating sequences of relatively complex frequency modulated (FM) sounds as the CS that either always (CS+) or never (CS−) predict the delayed US onset (Letzkus et al., 2011). Whereas auditory fear learning with simple sounds does not require neocortex, associative threat memories acquired through DTC depend upon higher order regions of the auditory cortex and, even more specifically, their descending projection to the BLA (Dalmay et al., 2019).

Inactivation studies establish the necessary involvement of descending corticoamygdalar (CAmy) projections in DTC without providing much insight into the nature or form of these neural changes. On the one hand, the key neural signatures of learning across a distributed hierarchy of brain areas can be reflected in the functional coupling between brain regions (Cambiaghi et al., 2016; Likhtik et al., 2013; Taub et al., 2018). On the other hand, establishing the essential neural changes underlying the acquisition and recall of discriminative threat memory has been greatly advanced through approaches that monitor and manipulate genetically targeted cell classes within the neocortex (Abs et al., 2018; Dalmay et al., 2019; Gillet et al., 2018; Letzkus et al., 2011), as well as specific glutamatergic (Belén Pardi et al., 2020), GABAergic (Schroeder et al., 2023), and cholinergic (Guo et al., 2019) input pathways to the auditory cortex. Either way, for relatively complex and naturalistic threat learning paradigms like DTC, where the neural substrates of threat memory require neocortical input to the BLA, recordings of unidentified cell types from one brain region at a time will be unlikely to reveal the nature and form of key underlying changes. Instead, progress on this front would require pulling the lens back to study changes in functional coupling between simultaneously recorded brain regions while also zooming in to pinpoint changes in particular cell types that support reorganization across distributed brain networks.

Here, we performed simultaneous recordings of single-unit spiking, local field potentials and ACh release in the BLA and higher-order auditory cortex (HO-AC) of awake, head-fixed mice during DTC. We used quantitative videographic measures of pupil and facial movement to index discriminative and generalized components of threat learning. Population measurements that indiscriminately pooled across neurons suggested enhanced CS discriminability in BLA but not HO-AC, yet optogenetically isolated recordings of CAmy projection neurons identified a subset of HO-AC neurons with a similar pattern of discriminative plasticity as observed in downstream BLA neurons. Asymmetrically enhanced functional coupling from the HO-AC to BLA (but not BLA to HO-AC) was observed at the end of a post-acquisition consolidation period and during threat memory recall. During acquisition, we also found that the sound-evoked ACh release was itself plastic and potentiated in BLA but not in HO-AC. Overall, our findings suggest that plasticity in the cholinergic and descending corticoamygdalar inputs facilitate the discriminative CS encoding in BLA upon threat learning.

## Results

### Sound-elicited facial movements and pupil dilation index DTC in head-fixed mice

Behavioral evidence of DTC in rodents is typically indexed via whole-body movements such as escape or freezing. Autonomic markers also provide a rapid and sensitive measure of DTC, with the additional advantage of lending themselves to head-fixed neural recording preparations (Weinberger and Diamond, 1987). Here, we performed Pavlovian auditory delay conditioning in head-fixed mice over three consecutive days alongside quantitative videographic measurements of the face and pupil. The first (habituation) and third (recall) sessions presented interleaved trials of five upward and downward frequency-modulated (FM) sweeps (**Figure 1A**). On Day 2 of DTC, a mild tail shock was initiated at the onset of the 5^th^ FM sweep. The FM sweep direction paired with tail shock (CS+) was counterbalanced across mice. In a separate cohort of Pseudo-conditioned mice, an equivalent number of tail shocks was presented during the intertrial interval, and was therefore not predictably related to either the upward or downward FM CS.

**Figure 1:**
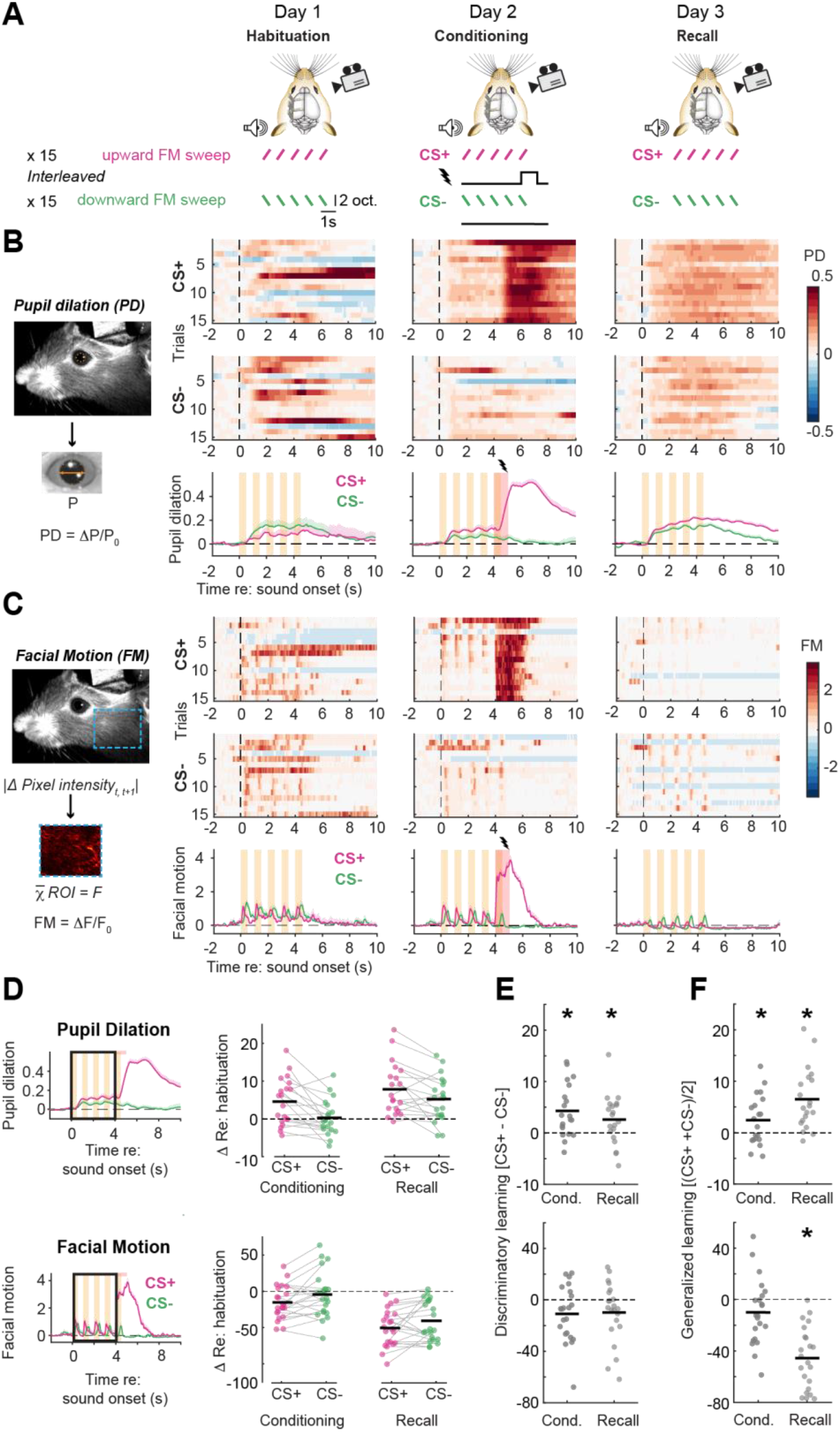
Pupil dilation and facial movements evoked by complex sounds index distinct timescales and conditioning specificity during auditory discriminative threat learning. **A**) Schematic illustrating the DTC protocol, where each of the three sessions are separated by 24 hours. In all the sessions, the mice are presented with 15 alternating presentations of a train of frequency modulated (FM) sweeps in upwards or downwards direction (conditioned stimuli, CS) during high-resolution facial videography. Upward sweeps are depicted as the CS+, though assignment of CS+ to sweep direction is counterbalanced across mice. **B**) *Left:* Pupil dilation (PD) in each trial is quantified as a fractional change in the pupil diameter (P) with respect to the mean pupil diameter in the 2s baseline before CS onset (ΔP/P_0_). *Right:* Fractional change in P for all CS presentations (*top* and *middle*) and mean pupil dilation across trials (*bottom*) for all three sessions in an example mouse. Vertical dashed lines denote onset of initial FM sweep, orange bars denote CS duration, and red bars denote the 1s shock. **C**) *Left:* Facial motion (FM) is computed at each time T as the absolute value of the difference in pixel intensities between consecutive frames (T, T+1) for each pixel and averaged over all the pixels within the region of interest (dashed blue rectangle). *Right:* Facial motion was expressed as a fractional change with respect to the mean facial motion in the 2s baseline before CS onset (ΔF/F_0_). Other plotting conventions match above. **D**) *Left:* Illustration of the area under the curve (AUC) quantification approach for pupil diameter and facial motion during the first 4 seconds of the CS presentation (black rectangle). *Right:* Difference in mean CS+ and CS− AUC for pupil and facial motion during Conditioning and Recall sessions relative to Habituation. Horizontal black bars indicate the mean. Pupil dilations were significantly larger for the CS+ and Recall Session: Repeated measures 2-way ANOVA, N = 20 mice; main effect for Stimulus [F = 14.51, p = 0.001], Session [F = 10.9, p = 0.004], no significant Session × Sound interaction [F = 1.72, p = 0.21]. Suppression of facial movements was greater for the CS+ during Recall (N = 22 mice): main effects for Sound [F = 9.61, p = 0.005], Session [F = 40.47, p < 2 × 10^−6^], no significant Session × Sound interaction [F = 0.02, p = 0.88]. **E**) Discriminatory changes reflect differences between the CS+ and CS−. Change in pupil diameter relative to Habituations session (shown above) for the CS− was subtracted from the CS+. *Top:* Discriminative changes in sound-evoked pupil dilations were larger in Conditioning than Recall (Repeated measures ANOVA, main effect for session F = 6.86, p = 0.003) but were significant in both sessions (one-sample t-tests with Bonferroni-Holm correction for multiple comparisons, Conditioning, p = 0.003; Recall, p = 0.025). *Bottom:* No significant discriminatory changes in facial movement were noted (Repeated measures ANOVA, main effect for session F = 2.28, p = 0.12; one-sample t-tests p = 0.06 for both after correction for multiple comparisons). **F**) Generalized changes reflect differences in Conditioning and Recall sessions that are CS non-specific. Change in pupil diameter relative to Habituations session was averaged for the CS+ and CS− stimuli. *Top:* Generalized increase in evoked pupil diameter was greater at Recall than Conditioning (Repeated measures ANOVA, main effect for session F = 14.47, p = 2 × 10^−5^), but were significant in both sessions (one-sample t-tests with Bonferroni-Holm correction for multiple comparisons, Conditioning, p = 0.04; Recall, p = 0.0002). *Bottom:* Generalized sound-evoked suppression of facial movements was significantly greater at Recall than Conditioning (Repeated measures ANOVA, main effect for session F = 39.68, p = 2 × 10^−10^), and was significant at Recall but not Conditioning (one-sample t-tests with Bonferroni-Holm correction for multiple comparisons, Conditioning, p = 0.08; Recall, p = 3 ×10^−8^).

Iso-luminous pupil dilations were elicited by the novel FM sweep stimuli during the Habituation session (**Figure 1B,** *left*) and also by the aversive unconditioned stimuli in the Conditioning session (**Figure 1B**, *center*). Pupil dilations also indexed discriminative learning, as evidenced by increased dilations beginning at the onset of the CS+ during the Conditioning and Recall sessions (**Figure 1B**, *right*) (Abs et al., 2018; Gehrlach et al., 2019; Oleson et al., 1972). We also noted that FM sweeps elicited rapid twitches of temporalis muscle that could be documented by measuring the motion energy within a region of interest positioned caudal to the vibrissa array (**Figure 1C**). Like isoluminous pupil dilations, facial motion was elicited by sound, by tail shock, and exhibited associative changes in response amplitude at the onset of the CS+ stimulus during the Recall session. Unlike pupil changes, facial motion tracked each individual FM sweep in the CS+ and CS− stimulus trains and was attenuated – rather than enhanced – at the onset of the CS+ (**Figure 1C,** *right*).

Conditioned behaviors can reflect generalized learning (non-discriminative changes to both the CS+ and CS−) and discriminative learning (larger changes in response to the CS+ than the CS−). To quantify the degree of generalized and discriminative learning in pupil dilations and facial movements, we quantified the overall response amplitude to the CS+ and CS− stimuli during the initial 4s of FM sweep trainings for the Habituation, Conditioning, and Recall sessions (**Figure 1D**). We found that pupil dilations reflected both significant discriminative (**Figure 1E)** and generalized (**Figure 1F)** learning on both the Conditioning and Recall sessions (statistical reporting provided in Figure Legends). By contrast, facial motion was not significantly changed during Conditioning and exhibited only generalized changes during Recall, confirming that autonomic conditioned responses are acquired more rapidly than motor conditioned responses (Weinberger and Diamond, 1987). Importantly, neither discriminative nor generalized changes in pupil diameter or facial movements were noted in Pseudo-Conditioned mice (**Supplemental Figure 1**). Taken together, these quantitative videographic measures demonstrate that pupil and facial movements index distinct timescales and forms of learning and confirm that DTC can be studied in head-fixed preparations that lend themselves to multiregional neurophysiological recording approaches.

### DTC increases the separability of neural population responses in BLA, not HO-AC

To characterize differences in the degree and form of associative plasticity in sensory cortex and the amygdala, we performed simultaneous single unit recordings from the HO-AC and BLA during DTC and Pseudo-conditioning (**Figure 2A**). HO-AC recordings targeted a lateral region of the auditory cortex labeled as AuV in the Allen Brain Institute Atlas or alternatively referred to either as A2 or SRAF in functional studies (Feigin et al., 2021; Narayanan et al., 2022; Romero et al., 2020; Stiebler et al., 1997). Post-mortem reconstructions confirmed that the vast majority of electrode positions aligned with AuV, though we cannot rule out the possibility that some electrode contacts might have been located in an even more lateral field, the temporal association area (TeA). Conservatively, we operationally define HO-AC to include AuV as well as the region of TeA adjacent to AuV.

**Figure 2:**
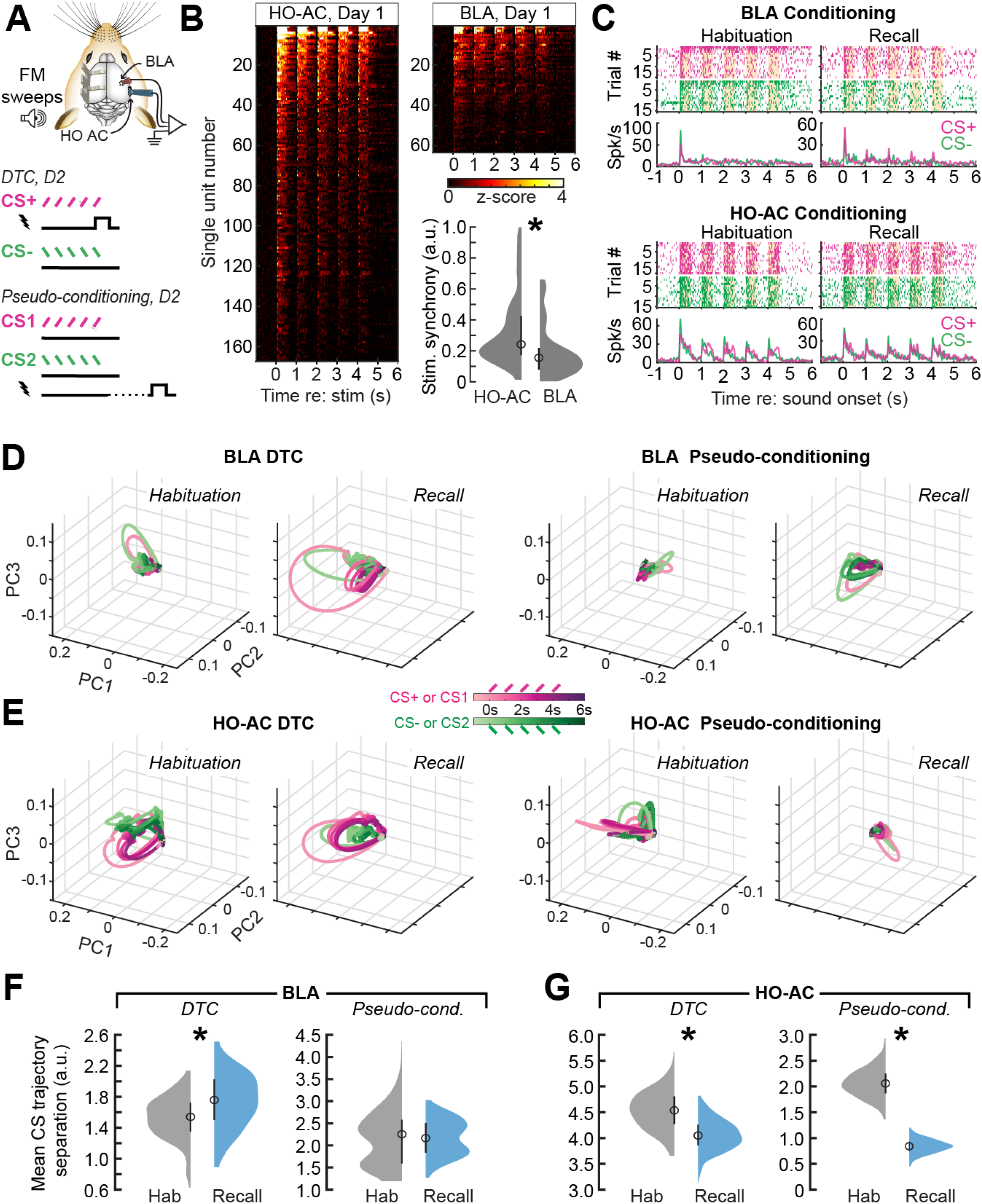
Complex sound representations become more separable after conditioning in BLA but become less separable over time in HO-AC. **A**) Extracellular single unit recordings were made with two 64-channel probes acutely positioned in the HO-AC and BLA on each day of the discriminative threat conditioning (DTC) or Pseudo-conditioning procedures. **B**) Neurograms showing 167 HO-AC and 63 BLA units recorded on the initial Habituation session to the five FM sweeps, 0.5s in duration, presented at 1Hz. Neurograms present one unit per row, where the spike rate is averaged over 30 upward and downward sweeps and expressed as a z-score. HO-AC units synchronized to the FM sweep train with significantly greater fidelity, as evidenced by a significantly greater amplitude of the Fourier transform at 1Hz (Wilcoxon rank-sum test, p value/Cliff’s delta = 1.41 × 10^−5^/-2.655). **C**) Rastergrams and peri-stimulus time histograms from four example units recorded on the Habituation and Recall sessions in the BLA (top) and HO-AC (bottom). **D**) BLA trial-averaged neural population responses throughout a 7s period surrounding the CS+ and CS− stimulus period is projected on a 3-dimensional space defined by the first three principal components (PCs). Stimulus trajectories expand and separate after DTC (left; N/n = 8/49 and 8/110 mice/units for Habituation and Recall, respectively) but remain relatively constricted and inseparable for both sessions in Pseudo-conditioned mice (right; 3/46 and 3/58 for Habituation and Recall, respectively). **E**) Same as above, but for HO-AC population responses during the Habituation and Recall sessions of DTC (N/n = 8/154 and 8/178 mice/units, respectively) and Pseudo-conditioning (N/n = 3/71 and 3/54, respectively). **F**) Euclidean distance between BLA CS+ and CS− population response trajectories averaged over the 5s CS duration (n = 500 bootstraps) was significantly increased in the Recall session compared to Habituation in DTC mice (*left*, unpaired t-test, p < 1 × 10^−10^; Cohen’s d = 0.68) but was not significantly changed in Pseudo-conditioned mice (*right;* unpaired t-test, p = 0.68; Cohen’s d = 0.03). **G**) Plotting conventions match above. HO-AC responses significantly habituate between the two recordings sessions, resulting in significantly less separable CS trajectories during the Recall session of both DTC (*left*, unpaired t-test, p < 1 × 10^−10^; Cohen’s d = −1.37) and Pseudo-conditioned mice (*right;* unpaired t-test, p < 1 × 10^−10^; Cohen’s d = −5.8).

During the initial Habituation session, upward and downward FM sweeps elicited responses from both regions, though the native, unconditioned sensory encoding fidelity was greater in the HO-AC, as evidenced by significantly greater synchronization of spike timing to each FM sweep within the 1Hz stimulus train (**Figure 2B**). After DTC, BLA units exhibited enhanced encoding of the CS+ but not CS− stimulus, by contrast to a representative HO-AC unit that showed equivalent responses to both stimuli in both recording sessions (**Figure 2C**).To measure changes in neural population-level stimulus discriminability before and after DTC, we visualized CS responses as trajectories in a reduced dimensionality space defined by the top three principal components (PCs) (Allsop et al., 2018; Dalmay et al., 2019). Before DTC, BLA population responses poorly differentiated between the train of upward and downward FM sweeps, reflecting stimulus adaptation and relatively poor synchronization. However, in the post-conditioning Recall session, BLA population responses displayed an elongated CS+ response trajectory that clearly diverged from the CS− (**Figure 2D**, *left*). By contrast, BLA response trajectories remained compressed and qualitatively indistinguishable for both Habituation and Recall sessions in Pseudo-Conditioned mice (**Figure 2D**, *right*). In an example population of HO-AC units, CS+ and CS− response trajectories appeared separable in both recordings sessions of DTC mice (**Figure 2E**, *left*) but became suppressed and poorly distinguished in the Recall session of a Pseudo-conditioned mouse (**Figure 2E**, *right*).

These observations were quantified by calculating the mean Euclidean distance between CS trajectories. In the BLA, CS+ and CS− responses were significantly more discriminable in Recall sessions compared to Habituation in DTC mice but not in pseudo-conditioned controls (**Figure 2F**). In HO-AC, neural population responses were robust and already distinct in the Habituation session, yielding larger CS trajectory separations than observed in BLA (**Figure 2G**). In Pseudo-conditioned mice, HO-AC units habituated to FM sweep stimuli by the third Recall session, resulting in compressed trajectories that were significantly less discriminable than the initial recordings. HO-AC responses in DTC mice exhibited a significant, albeit lesser degree of habituation found with Pseudo-conditioning, confirming other recent reports that DTC influences cortical population responses by counteracting habituation rather than increasing CS discriminability (Gillet et al., 2018; Wood et al., 2022).

### Robust associative plasticity in optogenetically targeted corticoamygdalar projection neurons

On a macroscopic scale, neural signatures of sensory associative learning are reflected in the strength and coherence of functional coupling between multiple brain areas (Cambiaghi et al., 2016; Herry and Johansen, 2014; Likhtik et al., 2013). These macroscopic changes are enabled by intrinsic and synaptic modifications in specific types of interneurons and projection neurons within local circuits (Letzkus et al., 2015; Pape and Pare, 2010). In this regard, blind recordings from single neurons in one brain area (e.g., Figure 2) can be both too precise and not precise enough. Focusing on learning-related changes in one brain area at a time provides no insight into potential changes in the strength or coherence between simultaneously recorded brain regions. On the other hand, collapsing across genetically or anatomically distinct classes of neurons can obscure highly localized plasticity within particular nodes of functional circuits. To address this point, we sought to both expand our focus to study dynamic changes in functional coupling between HO-AC and BLA during DTC while also narrowing our focus on corticoamygdalar (CAmy) projection neurons in the HO-AC that that innervate the BLA (**Figure 3A**).

**Figure 3:**
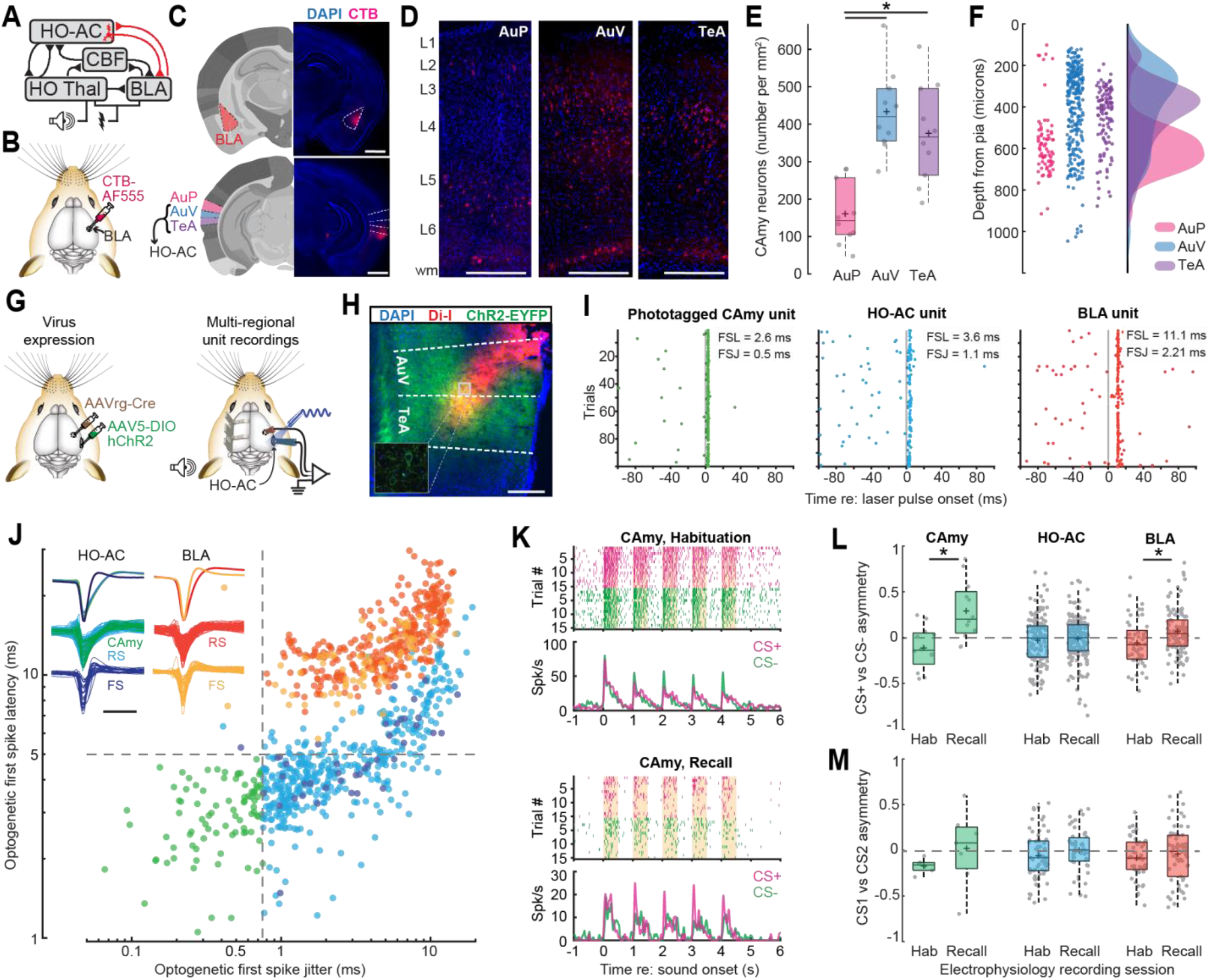
Associative plasticity in anatomically and optogenetically targeted HO-AC corticoamygdalar projection neurons resembles BLA neurons. **A**) Sensory representational plasticity reflecting the association of sound and shock is observed within the higher-order thalamus, cholinergic basal forebrain, auditory cortex, and BLA, but is less often studied at the level of isolated classes of corticofugal projection neurons or in the coherent activity between each brain area (illustrated in red). **B**) Cartoon illustrates injection of the fluorescent retrograde tracer CTB-AF555 into the BLA. **C**) Coronal sections depict the caudal portion of the BLA targeted for injection (top) as well as the primary region of AC (AuP), ventral AC (AuV), and temporal association area (TeA, bottom). Nomenclature and reference image at left are adapted from the Allen Institute for Brain Science. Here, HO-AC refers to regions denoted as AuV and TeA in the Allen Institute brain reference atlas. Fluorescence micrographs at right show CTB at the injection site as well as retrogradely labeled corticoamygadalar neurons (CAmy). Scale bar = 1mm. **D**) Density and laminar distribution of CTB+ CAmy neurons in AuP, AuV, and TeA. Scale bar = 0.25mm. **E)** Cell counts demonstrate that CAmy density is greater AuV and TeA compared to AuP (n = 10/5, slices/mice; Repeated measures ANOVA, F = 16.79, p = 2 × 10^−5^; post-hoc pairwise comparisons with Bonferroni-Holm correction for multiple comparisons, AuP vs AuV, p = 3 × 10^−5^; AuP vs TeA, p = 9 × 10^−4^; AuV vs TeA, p = 0.30). Box-and-whisker plots show median values in solid black lines, 25^th^ and 75^th^ percentiles and whiskers extending to the most extreme data points not considered outliers, + = mean. **F)** Distribution of CTB+ cells in AuP, AuV and TeA shown in *D* expressed as a function of distance from the pial surface. **G)** Cartoons illustrates strategy for selectively activating CAmy neurons via injection of a retrograde virus encoding cre-recombinase in the BLA and a cre-dependent virus encoding ChR2-EYFP in HO-AC (left). Multi-channel recording probes are positioned in BLA and HO-AC following a virus incubation period and brief pulses of 473nm light presented to the exposed cortical surface with a diode laser to activate ChR2+ CAmy neurons. **H)** Photomicrograph illustrates the Di-I coated silicon probe insertion trajectory in HO-AC relative to the approximate borders of AuV and TeA from the Allen Brain Institute reference atlas. Somata and neuropil of neurons transduced with both viruses express EYFP. Inset depicts a small region of interest photographed at higher magnification with a confocal microscope to illustrate somatic expression of ChR2-EYFP. Scale bar = 0.25 mm. **I**) Spike rasters from a photo-tagged CAmy unit and RS single units in HO-ACtx and BLA in response to the 1ms laser pulse stimulation. Gray vertical line denotes onset of 1 ms laser pulse. FSL = first spike latency. FSJ = first spike jitter. **J**) *Inset:* HO-AC and BLA single units were classified as regular spiking (RS) or fast spiking (FS) (trough-to-peak delay >= 0.6 ms or < 0.6 ms, respectively). Mean ± SEM; waveform shapes shown on top row; waveforms from all units shown in bottom two rows. Scale bar = 1ms. CAmy units (green) were operationally defined as HO-AC RS units with a low first spike latency (< 5 ms, dashed horizontal line) and a low first spike jitter (< 0.75 ms, dashed vertical line) in response to a 1 ms laser pulse stimulation. All other RS and FS units in HO-AC and BLA are also plotted for comparison. **K**) Rastergrams and peri-stimulus time histograms from two example CAmy units recorded on the Habituation and Recall sessions. **L**) Discriminative plasticity from sound-responsive units in 8 mice that underwent DTC using an asymmetry index ((CS+ - CS−) / (CS+ + CS−), where positive values reflect a greater response to the CS+, negative values to the CS− and a value of zero reflects an equivalent response to both stimuli. CS− evoked responses were significantly more biased towards the CS+ in the Recall session compared to Habituation in BLA RS units (n = 49/110 Habituation/Recall; unpaired t-test, p = 0.003, Cohen’s d = 0.51) and optogenetically phototagged HO-AC CAmy units (n = 12/12 Habituation/Recall; p = 0.002, Cohen’s d = 1.44), but not HO-AC in RS units that were not identified as CAmy units (n = 142/166 Habituation/Recall; p = 0.59, Cohen’s d = 0.06). **M**) Discriminative plasticity from sound-responsive units in 3 mice that underwent Pseudo-Conditioning with the same analysis described above. CS− evoked responses did not show a significant difference in bias in BLA RS units (n = 46/58 Habituation/Recall; unpaired t-test, p = 0.43, Cohen’s d = 0.16), optogenetically phototagged HO-AC CAmy units (n = 6/7 Habituation/Recall; p = 0.29, Cohen’s d = 0.62), or HO-AC RS units not identified as CAmy units (n = 65/47 Habituation/Recall; p = 0.22, Cohen’s d = 0.24).

To record from isolated CAmy neurons, we first sought to determine their laminar and areal distribution within the auditory cortex. This was accomplished by injecting a retrograde tracer, CTB, into the BLA (**Figure 3B**) and documenting their abundance and cortical depth in primary regions of AC (AuP), the ventral AC – (AuV), and in temporal association cortex lateral to AC (TeA) (**Figure 3C-D**). We observed approximately twice as many CTB-labeled CAmy neurons in AuV and TeA as in AuP (**Figure 3E**) and noted that CAmy neurons were distribute across the cortical column – approximately from layer 2 to layer 5 – in AuV and TeA but were mostly restricted to layer 5 in AuP (**Figure 3F**). These anatomical findings affirm that more lateral HO-AC regions are more strongly connected with the BLA.

To record from CAmy units, we used an intersectional virus strategy to inject a retro-Cre virus in BLA and a cre-dependent virus in HO-AC to limit the expression of channelrhodopsin (ChR2) to CAmy neurons (**Figure 3G**). After allowing the virus several weeks to incubate, we then made translaminar recordings from HO-AC (**Figure 3H**) and quantified the latency and temporal jitter of spikes evoked by a 1ms pulse of blue light to the cortical surface (**Figure 3I**). Following a conservative approach used in previous studies (Guo et al., 2019; Nieh et al., 2015; Williamson and Polley, 2019), optogenetically tagged CAmy units were distinguished from indirectly activated HO-AC and BLA units based on the strength of evoked spiking (at least 5 SD above baseline), the shorter latency of direct versus polysynaptic activation (less than 5ms from laser onset), and highly stereotyped spike timing across trials (less than 0.75ms of jitter; **Figure 3J**).

Like other HO-AC single units recorded during DTC, example CAmy units showed robust, non-adapting responses to FM sweeps during the initial recording session and more suppressed, habituated responses on the Day 3 Recall session (**Figure 3K**). However, CAmy units also exhibited characteristics seen only in BLA units, in that CS+ responses were enhanced relative to CS− on the Recall session. We quantified these changes across regular spiking units in each brain region with an asymmetry index, where positive values indicated a response bias towards the CS+, negative values a bias towards the CS−, and a value of zero reflecting balanced spike rates to upward and downward FM sweeps (**Figure 3L**). CAmy responses were significantly biased towards the CS+ during Recall compared to Habituation, matching the relationship observed in BLA units but in contrast to neighboring HO-AC units, which did not show a significant CS response bias in either recording session. As a negative control, significant CS bias was not observed in BLA, HO-AC, or CAmy units in Pseudo-conditioned mice (**Figure 3M**). Finally, CS+ response bias was also not observed in fast-spiking units from either brain region with DTC or Pseudo-conditioning, which further underscores the cell type-specific expression of associative plasticity in both brain regions (**Supplemental Figure 2**) (Gillet et al., 2018; Guo et al., 2019; Krabbe et al., 2018).

### Enhanced corticoamygdalar-evoked local network responses in BLA after DTC

Prior work has shown that selective optogenetic inactivation of auditory CAmy axons blocks the behavioral retrieval of threat memory, suggesting that auditory corticofugal projection neurons transmit critical information to the amygdala, particularly for complex auditory CS stimuli (Dalmay et al., 2019). In a similar vein, plasticity of auditory corticofugal synapses in the posterior striatum is necessary for perceptual learning in an operant auditory frequency discrimination task, where a signature of this plasticity can be studied *in vivo* via enhanced LFP amplitude in the striatum elicited by optogenetic activation of corticostriatal projection neurons (Xiong et al., 2015). To test the hypothesis that a similar enhancement of a CAmy-evoked LFP would be evident in the BLA following DTC, we measured the BLA LFP response to a brief (1ms) optogenetic activation of CAmy projection neurons at varying laser powers (**Figure 4A**).

**Figure 4:**
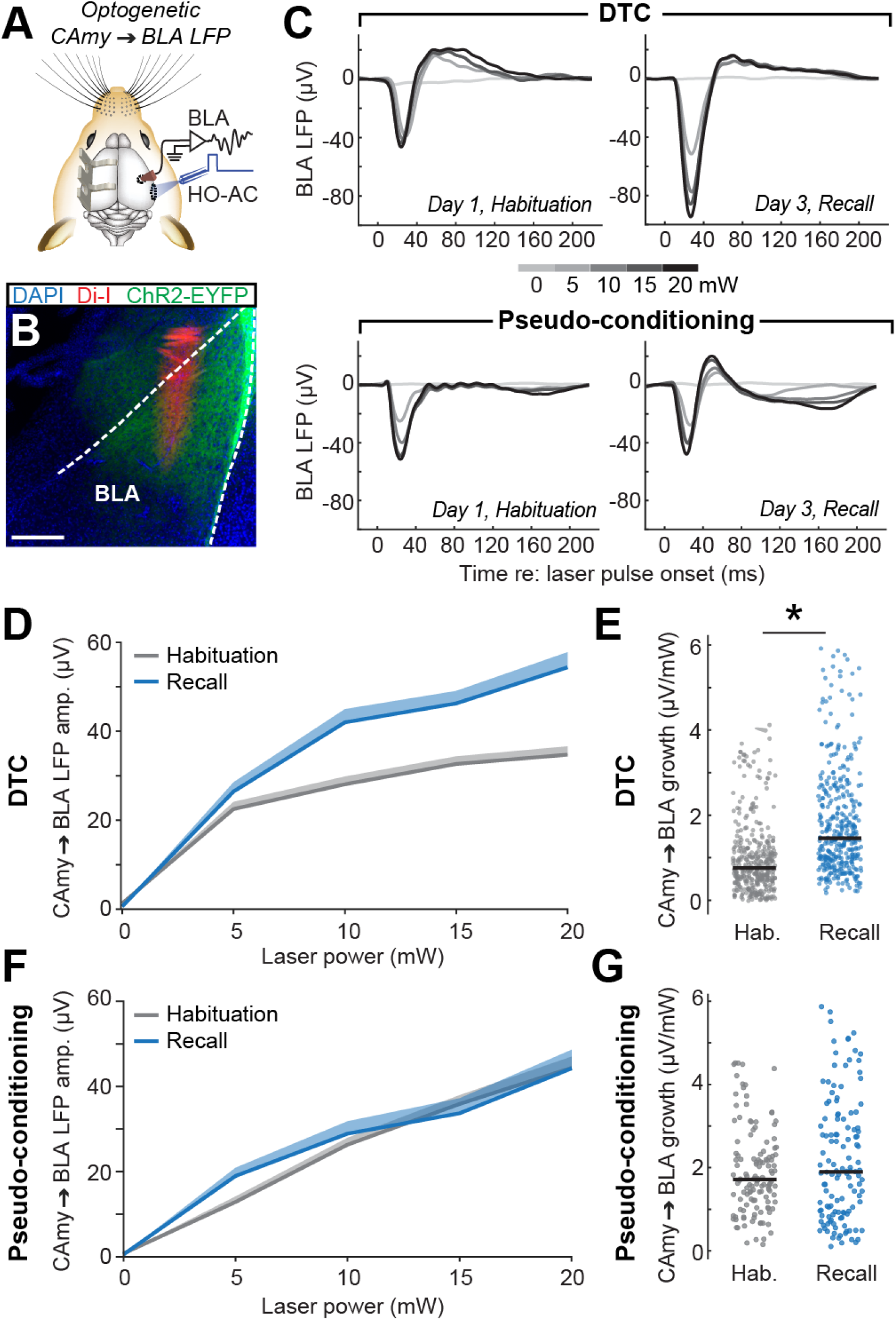
Potentiation of Corticoamygdalar-evoked BLA activity after discriminative threat conditioning. **A**) Cartoon illustrating the protocol for BLA local field potential (LFP) recordings during bulk optogenetic stimulation of HO-AC CAmy neurons. **B**) Photomicrograph illustrates the Di-I coated silicon probe insertion trajectory in BLA and ChR2-EYFP+ cortical axon terminals. Dashed lines demarcate approximate BLA border based on Allen Brain Institute Reference Atlas. Scale bar = 0.25 mm. **C**) Optogenetically evoked LFPs in the BLA of an example DTC mouse (top) and Pseudo-conditioned mouse (bottom). **D**) Mean ± SEM LFP amplitude as a function of laser power in the example DTC mouse during Habituation and Recall. **E**) Slopes from the light-evoked response growth functions for all channels in DTC mice (n = 384/6, channels/mice) indicate enhanced response growth during recall compared to habituation. Horizontal black bars indicate the median. Asterisks indicate statistical significance with Wilcoxon rank-sum test (p value/Cliff’s delta = 2 × 10^−30^/0.48). **F**) As per *D*, but in an example Pseudo-conditioned control mouse. **G**) As per *E*, but in Pseudo-conditioned controls (n = 128/2, channels/mice). Slopes from the light-evoked response growth functions for all channels indicate no significant change in the response growth during recall compared to habituation; Wilcoxon rank-sum test (p value/Cliff’s delta = 0.46/0.05).

As expected, axon terminal expression of ChR2-EYFP in the BLA was robust (**Figure 4B**), and optogenetic activation of CAmy cell bodies in HO-AC elicited a monotonic increase in BLA LFP amplitude with increasing laser power (**Figure 4C**). In mice undergoing DTC, CAmy-evoked LFP amplitude was significantly greater during the Recall session than Habituation session (**Figure 4D**), yielding a significantly greater growth slope across laser powers (**Figure 4E**). No significant changes in CAmy-evoked BLA LFP amplitude or growth slopes were noted between the Habituation and Recall sessions of mice that underwent Pseudo-conditioning (**Figure 4F-G**). Importantly, the optogenetic activation protocol was performed just prior to the interleaved presentation of upward and downward FM sweeps on the Habituation and Recall sessions, thus highlighting a stabilized potentiation of CAmy efferents that persisted for at least 24 hours following DTC.

### Asymmetric potentiation of corticoamygdalar –not amygdalocortical – inputs during threat memory recall

To test the hypothesis that HO-AC inputs to the BLA are enhanced during the recall of threat memory during naturally occurring patterns of neural activity, we measured the BLA LFP triggered by HO-AC spiking during the CS presentation period (**Figure 5A**). The spike-triggered LFP indexes transient changes in the strength and timing of information flow between brain regions (Einevoll et al., 2013) and has been used in prior studies to identify an enhanced functional coupling between the amygdala and pre-frontal cortex (Taub et al., 2018), as well as basal forebrain cholinergic neurons and auditory cortex during trace auditory fear conditioning (Guo et al., 2019) and auditory operant learning (Laszlovszky et al., 2020). To mitigate noise from other neural sources or other spikes occurring at short intervals, we used a linear deconvolution method rather than simply calculating the spike-triggered average (Ehinger and Dimigen, 2019).

**Figure 5:**
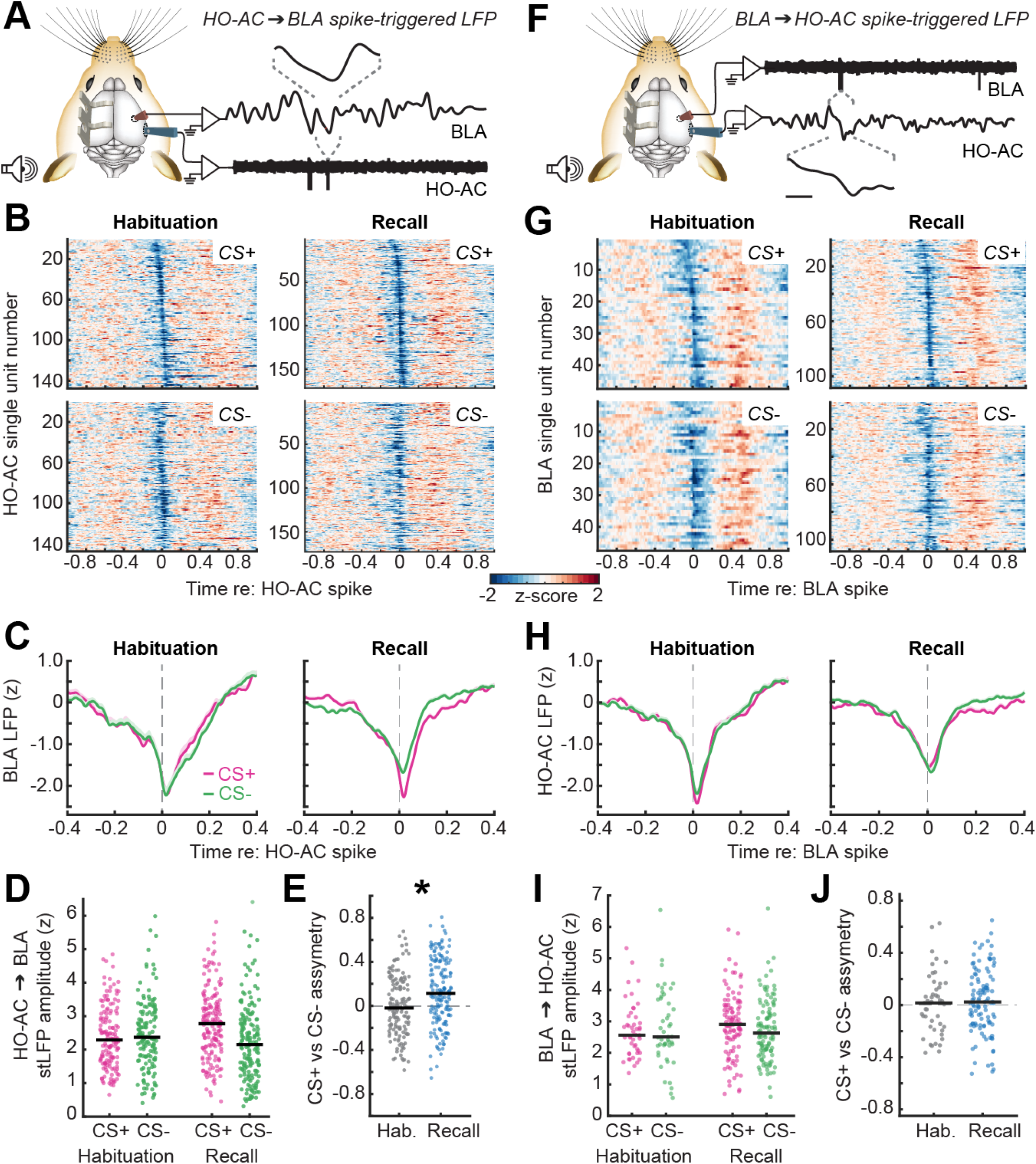
Enhanced functional coupling from HO-AC to BLA - but not BLA to HO-AC - during threat memory recall. **A**) Schematic illustrating the quantification of BLA LFPs triggered by HO-AC single unit spikes. Linear deconvolution by time expansion is used to estimate the spike-triggered LFP (stLFP). **B**) Estimated HO-AC to BLA stLFPs computed during the CS+ (*top*) and *CS−(bottom*) expressed as a z-score relative to pre-stimulus baseline and averaged across all recording channels in BLA. **C**) Mean ± SEM HO-AC to BLA stLFP demonstrates a downward deflection of the BLA shortly following HO-AC spikes that is selectively enhanced during the CS+ period in the Recall session. **D**) BLA stLFP amplitude for each HO-AC RS unit during each CS presentation on Habituation and Recall sessions (N/n = 8/147, 8/171 mice/units for Habituation and Recall respectively). Horizontal black bars indicate the mean. BLA stLFP is significantly and specifically elevated during CS+ stimuli after DTC: Mixed model ANOVA with Session as a factor and Sound as a repeated measure, main effect for Session [F = 0.86, p = 0.35], main effect for Sound [F = 8.88, p = 0.003], Session × Sound interaction term [F = 15.43, P = 0.0001]. **E**) Discriminative plasticity in the HO-AC to BLA stLFP for each unit can be expressed as an asymmetry index ([CS+ – CS−] / [CS+ + CS−] where positive values reflect a greater response to the CS+, negative values to the CS− and a value of zero denotes an equivalent response. The asymmetry index was significantly greater than zero in the Recall session (one-sample t-test, p = 8 × 10^−7^; Cohen’s d = 0.41) and was significantly more positive than the Habituation session (unpaired t-test, p value/Cohen’s d = 0.0002/0.44). Horizontal black bars indicate the mean. **F-H**) As per *A-C*, but for the HO-AC LFP triggered by spikes in individual BLA units. **I**) Plotting conventions match *D*. HO-AC stLFP amplitude for each BLA unit (N/n = 8/47, 8/108 mice/units for Habituation and Recall respectively). Horizontal black bars indicate the mean. No significant changes were observed: Mixed model ANOVA with Session as a factor and Sound as a repeated measure, main effect for Session [F = 0.52, p = 0.47], main effect for Sound [F = 0.28, p = 0.6], Session × Sound interaction term [F = 1.57, p = 0.21]. **J**) Plotting conventions match *E*. BLA to HO-AC stLFP amplitude was not significantly biased towards the CS+ during Recall (one-sample t-test, p = 0.62, Cohen’s d = 0.12) and was not significantly different than in Recall compared to Habituation (Unpaired t-test, p value/Cohen’s d = 1.29/0.08). Horizontal black bars indicate the mean.

We noted that HO-AC spikes were associated with negative deflections of the BLA that peaked 5-10ms after the cortical spike (**Figure 5B**). The spike-triggered LFP was equivalent for CS+ and CS− stimuli during the initial Habituation session but was discriminatively enhanced during the CS+ presentation period in the Recall session (**Figure 5C-E**). Importantly, the sound-evoked LFP amplitude did not differ between CS+ and CS− stimuli in HO-AC or BLA (**Supplemental Figure 3**), confirming that associative plasticity in the spike-triggered LFP during the CS presentation period reflects an enhanced functional connection between the HO-AC and BLA and cannot be solely explained by a bottom-up change in the sound-evoked LFP.

BLA neurons receive direct anatomical inputs from the HO-AC but also directly project to the HO-AC (Tasaka et al., 2020; Tsukano et al., 2019). To determine whether enhanced functional connectivity between the HO-AC and BLA during threat memory is bi-directional or asymmetric, we also calculated the spike-triggered LFP from BLA to HO-AC (**Figure 5F**). We also noted that negative deflections in cortical LFPs peaked several millisecond following BLA spikes, confirming corticopetal functional connectivity (**Figure 5G**). However, we did not observe a systematic difference in the spike-triggered LFP amplitude between CS+ and CS− stimuli in either recording session (**Figure 5H-J**). Further, CS− specific changes in the spike-triggered LFP were not observed for either direction in mice that underwent the Pseudo-conditioning protocol (**Supplemental Figure 4**), underscoring that enhanced functional connectivity was specific to the CS+ during DTC Recall and only for descending corticofugal projections.

### Discriminative changes in amygdalar ACh release during threat acquisition

Direct recordings from neuromodulatory inputs to the HO-AC and BLA have shown that they function like teaching signals, on account of their short-latency phasic responses to auditory stimuli that are rapidly and discriminatively rescaled when they are predictive of aversive stimuli (Crouse et al., 2020; Guo et al., 2019; Robert et al., 2021; Schroeder et al., 2023). This raises the possibility that CS sounds might elicit endogenous ACh release in both brain structures and that the amplitude of ACh release could be discriminatively modified early in the DTC process, even during the Acquisition session. While tail shock electrically interfered with our ability to make single unit recordings during the Acquisition session, ACh release can be measured optically, which we reasoned would allow us to determine whether discriminative plasticity in cholinergic inputs to both HO-AC and BLA were linked to discriminative plasticity in spiking responses and functional connectivity described above, which were measured during the subsequent threat memory consolidation and recall period.

To measure endogenous ACh release dynamics throughout all stages of DTC, we expressed the genetically encoded ACh fluorescent sensor, GRAB_ACh_3.0 (ACh3.0; **Figure 6A**; Jing et al., 2020), and monitored fluorescence dynamics in the BLA (**Figure 6B**) and HO-AC (**Figure 6C**) simultaneously with dual optic fiber implants. To leverage the advantages of fiber photometry for stable long-term recordings and to capture ACh dynamics with greater sensitivity during acquisition, we extended the Habituation phase of the DTC procedure to two days and the Conditioning phase to three days. A final post-conditioning session on day 6 provided an assay for threat memory recall. On the first session, we noted robust sound-evoked ACh release in both BLA (**Figure 6D**) and HO-AC (**Figure 6E**). As observed previously, sound-evoked cholinergic responses were steeply reduced on Day 2 of the Habitation session, particularly in HO-AC, reflecting strong habituation to stimulus novelty (**Figure 6E-F**) (Robert et al., 2021). Across the three Conditioning sessions, the 1s tail shock stimulus combined with the fifth CS+ stimulus to produce an even greater surge in local ACh release. For both the auditory CS and the tail shock, ACh release appeared more phasic in BLA and more protracted in HO-AC, perhaps reflecting differences in acetylcholinesterase levels in each brain region.

**Figure 6:**
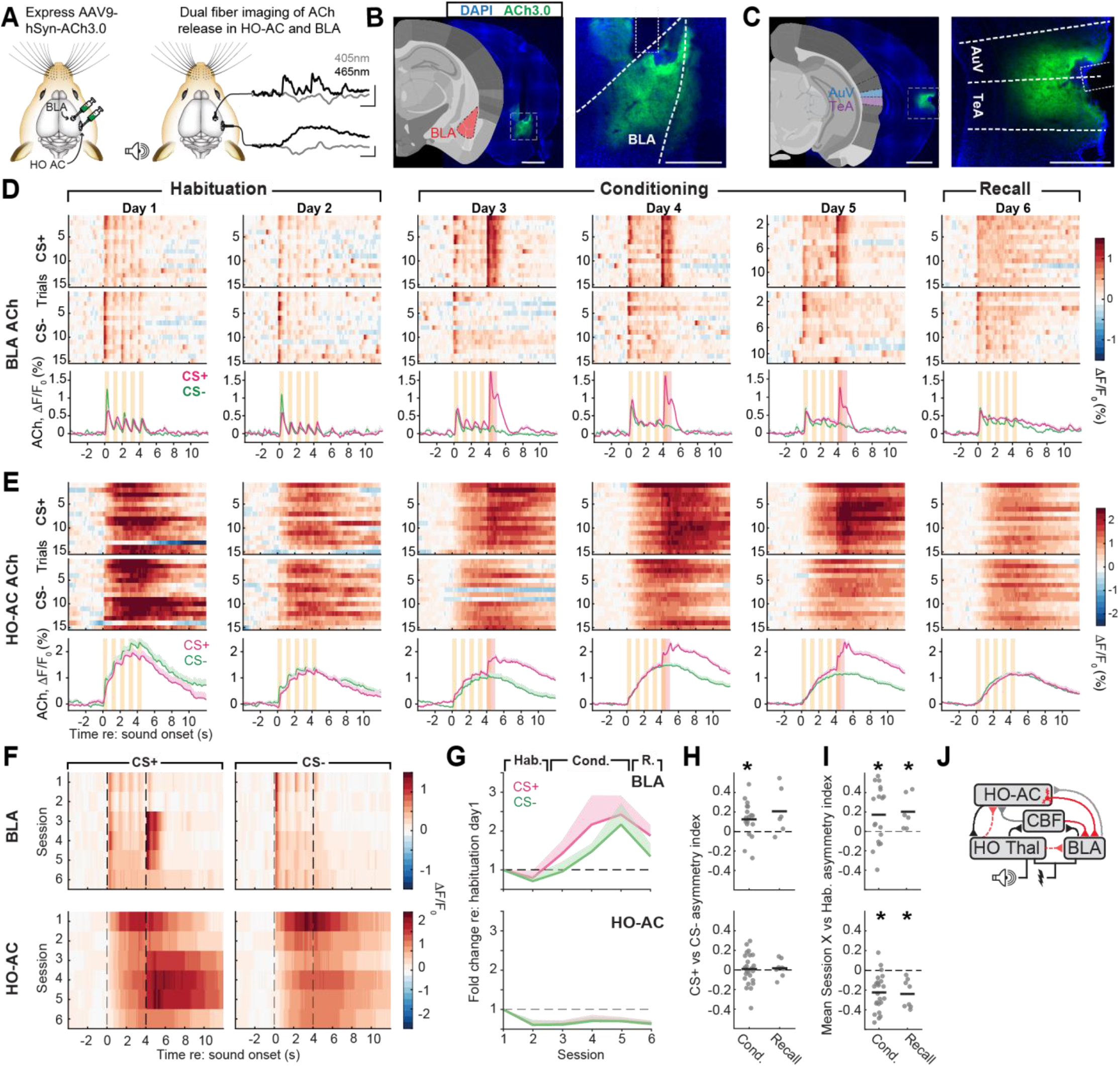
Discriminative and generalized changes in sound-evoked acetylcholine release begin during threat conditioning. **A)** *Left:* Schematic illustrating viral expression of the genetically encoded fluorescent ACh sensor, ACh3.0. *Right:* Simultaneous fiber-based bulk fluorescence measurements in the BLA and HO-AC at the ACh3.0 excitation wavelength (465nm) and a control wavelength (405) that is not sensitive to ACh release. Vertical and horizontal bars represent 1% ΔF and 2s, respectively. **B)** Coronal sections depict the caudal portion of the BLA targeted for injection (left) Nomenclature and reference image are adapted from the Allen Institute for Brain Science. Fluorescence micrographs show ACh3.0 expression and estimated position of the optic fiber (dotted white line). Scale bars are 1mm at left and 0.5mm at right. **C)** As per *B*, but for HO-AC. AuV and TeA are naming conventions from the Allen Institute for Brain Science reference atlas denoting the ventral auditory cortex and temporal association area, respectively. Scale bars are 1mm at left and 0.5mm at right. **D)** The fractional change in fluorescence provides a measure of endogenous ACh release in BLA elicited by each FM sweep (yellow rectangle) and tail shock (red rectangle) in an example mouse. For fiber imaging experiments, the Habituation phase is extended to two days and the Conditioning phase to three days. **E)** As per *D*, but for HO-AC recorded simultaneously with the BLA probe in the same mouse. **F)** Mean trial-averaged fractional change for each recording session across six dual implant mice. Dashed vertical lines denote the 4s CS period prior to onset of the 5^th^ FM sweep and tail shock used to calculate the CS+ and CS− response values. **G)** Mean ± SEM fold change in CS+ and CS− evoked activity during the initial 4s stimulus period expressed relative to the first Habituation session in six dual implant mice. In BLA (*top*), sound-evoked cholinergic responses increase throughout Conditioning and Recall but do not systematically differ by CS type: 2-way repeated measures ANOVA, main effect for Session [F = 5.08, p = 0.002], main effect for Sound [F = 3.11, p = 0.14], Session × Sound interaction term [F = 0.7, p = 0.63]. In HO-AC (*bottom*), sound-evoked cholinergic responses decrease throughout Conditioning and Recall but do not systematically differ by CS type: 2-way repeated measures ANOVA, main effect for Session [F = 4.5, p = 0.003], main effect for Sound [F = 0.07, p = 0.8], Session × Sound interaction term [F = 0.03, p = 0.99]. **H)** Differences in evoked responses by the CS+ and CS− are calculated from the response to each stimulus for a given Conditioning or Recall session relative to the mean of the two Habituation sessions. Discriminative plasticity is computed as an asymmetry index ([CS+ – CS−] / [CS+ + CS−]) where a positive value denotes a greater response to the CS+. Circles denote the trial-averaged mean for a single session from a single mouse. Horizontal black lines denote sample mean. In BLA (*top*), cholinergic responses were significantly greater for the CS+ than CS− during the Conditioning sessions (one-sample t-test relative to a population mean of zero, p = 0.01, Cohen’s d = 0.68, n = 18) and was marginally greater during the Recall session (p = 0.09, Cohen’s d = 0.87, n = 6). Significant discriminatory changes were not observed in HO-AC (*bottom;* p > 0.58 and Cohen’s d < 0.21 for both Conditioning and Recall, n = 24 and 8 respectively). **I)** As per H, but CS− evoked responses were averaged across the CS+ and CS− to provide an measure of generalized changes over threat memory acquisition and retrieval. Generalized plasticity is computed as an asymmetry index ([Session X – Habituation_mean_] / [Session X + Habituation_mean_]) where a positive value denotes the mean CS response is greater than Habituation. In BLA (*top*), CS− evoked cholinergic responses were significantly increased relative to Habituation in both Conditioning and Recall (p < 0.04 and Cohen’s d > 0.58 for both time points). In HO-AC (*bottom*), CS− evoked cholinergic responses were significantly reduced relative to Habituation in both Conditioning and Recall (p < 0.002 and |Cohen’s d| > 1.23 for both time points). **J)** Summary diagram illustrating afferent pathways to the BLA shown here to be discriminatively modified during the acquisition, consolidation, or recall of threat memory (solid red). Other projection pathways shown in other work to be discriminatively modified during DTC are shown in dashed red. Gray lines denote pathways studied here that did not exhibit discriminative change. Black denotes pathways that have not yet been investigated in the context of DTC.

To capture learning-associated changes in ACh release while excluding time periods associated with tail shock and resultant changes in movement and arousal, we quantified the integrated fluorescence response during the initial 4s of the auditory cue period (**Figure 6F**, dashed lines). We noted a striking divergence in sound-evoked ACh release between brain regions. In BLA, CS− evoked responses increased across Conditioning and Recall sessions, particularly for the CS+ (**Figure 6G**, *top*), thus paralleling the BLA neural population responses measured during Recall (Figure 2D and 2F). By contrast, HO-AC ACh release strongly habituated over time for both the CS+ and CS− (**Figure 6G**, *bottom*), again paralleling the net reduction in stimulus discriminability noted in HO-AC population responses (Figure 2E and 2G). We found significant discriminatory plasticity – enhanced ACh release for the CS+ relative to the CS− – during Conditioning in the BLA (**Figure 6H**, *top*), paralleling the conditioned changes noted in pupil dilations (Figure 1E), but no discriminative changes in HO-AC. Although opposite in sign – increased in BLA, decreased in HO-AC – CS− evoked ACh release showed significant generalized changes during the Conditioning sessions for both brain regions (**Figure 6I**).

## Discussion

Here, we studied DTC in head-fixed mice using relatively complex, naturalistic sounds as conditioned stimuli and sound-evoked changes in pupil dilation and facial twitches as behavioral indices of both specific and generalized threat memory (Figure 1). Simultaneous multi-regional recordings identified enhanced CS discriminability in BLA population responses, whereas CS population responses in HO-AC habituated over time, making the cortical representations less discriminable (Figure 2). At a single-unit level, we noted a cell-type specific potentiation in the CS+-evoked responses in photo-tagged CAmy units that was not observed in unidentified neighboring HO-AC units but was comparable to BLA units (Figure 3). To investigate the dynamics in functional connectivity between brain regions, we optogenetically activated CAmy neurons while recording LFP responses in BLA or alternatively used the natural spiking of BLA and HO-AC units as the LFP trigger. We found that direct, bulk activation of CAmy projection neurons prior to recall elicited potentiated network level responses in BLA (Figure 4). HO-AC spike-triggered LFPs in the BLA were also significantly potentiated during CS+ stimulus presentation at recall, whereas BLA spike-triggered cortical LFPs showed no change (Figure 5). Taken together, our findings show that threat memories are encoded by BLA ensembles and reflect a selective potentiation of descending CAmy inputs, without necessarily reflecting a gross reorganization of non-specific HO-AC ensemble responses. As a final point, pupil dilations indexed significant discriminative learning during threat memory acquisition, which was paralleled by elevated CS+-evoked ACh release in the BLA during the Acquisition period. Further, divergent population-level neural reorganization in the Recall session – indiscriminate CS habituation in HO-AC but discriminative CS enhancement in BLA – was paralleled by suppressed sound-evoked ACh release in HO-AC and enhanced sound-evoked ACh release in BLA. These findings suggest that reorganized cholinergic inputs may guide – rather than simply enable – generalized and discriminative changes in neural sound processing in both brain regions (**Figure 6J**), though the strong of that hypothesis awaits future studies that would employ targeted inactivation protocols to test the necessity of each afferent input for BLA plasticity and behavioral memory strength.

### Differences in the degree, form, and specificity of plasticity underlying auditory threat memory in BLA and auditory cortex

Across animal models and conditioning protocols, there is strong overall evidence for a rapid and persistent reorganization of BLA responses to enhance the salience of sounds that predict aversive reinforcement (Janak and Tye, 2015; LeDoux, 2007). Selective enhancement of CS+ representations following DTC been reported in the auditory cortex (Weinberger, 2004), though cortical reorganization is less consistent overall than BLA and depends – as we have shown here – on the cortical cell type and auditory cortex region (Abs et al., 2018; Dalmay et al., 2019), the degree of generalized versus specific fear learning (Aizenberg and Geffen, 2013; Wood et al., 2022), the use of complex auditory CS stimuli or more complex conditioning protocols (Dalmay et al., 2019; Gillet et al., 2018; Guo et al., 2019), and has been interpreted as reflecting attentive processing of threatening stimuli rather than their short latency encoding (Quirk et al., 1997).

We noted a selective enhancement of the CS+ representation in BLA population responses and regular spiking unit firing rates but not in HO-AC population responses or single unit firing rates. Discriminative plasticity in CAmy units, by contrast, were more akin to BLA units than to neighboring units in HO-AC, in that they also exhibited a selective enhancement of CS+ response. These findings can be explained by a dual-stream model, which purports that the auditory thalamus and cortex feature intermingled functional populations of highly plastic neurons that reflect the learned significance of environmental sounds (e.g., CAmy projection neurons) alongside other populations that are optimized for stability to encode environmental stimuli based on their physical features and overall novelty independent of fear associations (Gründemann, 2021; Leppla et al., 2022). Alternatively, unidentified regular- and fast-spiking units that on average did not exhibit discriminative enhancement of the CS could nevertheless encode associative threat memory at more remote time point than the next-day Recall session used here (Cambiaghi et al., 2016; Concina et al., 2022; Yang et al., 2016). A third possibility is that most HO-AC neurons do encode the discriminative threat memory at the time scale studied here, but the representation of the memory is not based in overall changes in firing rate but instead in the stability of neurons that are functionally connected into CS+ and CS− ensembles (Dalmay et al., 2019; Grewe et al., 2017; Taylor et al., 2021; Wood et al., 2022).

### Inter-regional functional coupling and asymmetric potentiation in corticofugal plasticity

The BLA and HO-AC are reciprocally interconnected, where the HO-AC both sends and receives approximately three time more input with the BLA than AuP, as shown here and in prior work (Hintiryan et al., 2021; LeDoux et al., 1991; Romanski and Ledoux, 1993; Tsukano et al., 2019; Yang et al., 2016). We used the spike-triggered LFP to demonstrate that the reciprocal anatomical connectivity between HO-AC and BLA is mirrored by reciprocal functional connectivity, such that a spike in either region was associated with the maximal negativity in the LFP 5-10ms later, the temporal lag suggesting that the major contributor is the inter-area communication rather than shared common inputs. Previously spike-triggered LFPs have been used to study the coupling of amygdala spikes to prefrontal cortex LFPs during threat conditioning (Taub et al., 2018), and cholinergic basal forebrain spikes to auditory cortex LFPs during auditory trace fear conditioning (Guo et al., 2019), and operant learning (Laszlovszky et al., 2020). The directional coordination between the output (spikes) of one region and the input (local field potentials) of another brain region is thought to facilitate learning and memory encoding of salient information by reducing inter-trial variability and increasing postsynaptic excitability, thereby allowing for an efficient information transfer (Taub et al., 2018).

Sensory corticofugal neurons innervate far-flung targets in the forebrain, midbrain, and brainstem and their plasticity can shape real-time processing and guide long-term reorganization of their downstream subcortical targets (Asokan et al., 2018; Gao and Suga, 1998; Liu et al., 2016; Zingg et al., 2017). Despite the symmetry in the native functional connectivity between HO-AC and BLA, only the CAmy projection neurons and descending functional coupling assays exhibited discriminative plasticity. The asymmetric potentiation in CAmy influence on BLA ensembles reinforces inactivation studies showing the necessary involvement HO-AC CAmy projections in the recall of short-term threat memory with complex sounds (Dalmay et al., 2019), in the recall of remote auditory threat memories (Cambiaghi et al., 2016), and in the re-acquisition of additional auditory threat associations (Concina et al., 2022). Future work is needed to address whether the asymmetric CS+ response potentiation in the CAmy neurons is distinct from other projection neuron types across the cortical column. One possibility is that compared to other pyramidal neuron types, the apical dendrites of CAMy projection neurons are preferentially targeted by layer 1 interneurons or long-range afferents from higher-order regions of the auditory thalamus and zona incerta that are also concentrated in layer 1 and all exhibit strong discriminative enhancement of the CS+ representation(Abs et al., 2018; Belén Pardi et al., 2020; Letzkus et al., 2011; Schroeder et al., 2023).

### Cholinergic modulation in BLA and HO-AC

Complex sounds even with no learned relevance evoked ACh release in both HO-AC and BLA during the Habituation session. Over the subsequent days of conditioning, sound-evoked ACh release was indiscriminately reduced in HO-AC but selectively enhanced for the CS+ in BLA. ACh acts on BLA principal neurons via muscarinic receptors to prolong depolarization, promote long-term plasticity of CAMy and local synapses and enable the acquisition of fear memories (Crimmins et al., 2022; Jiang et al., 2016; Kellis et al., 2020; Unal et al., 2015). In this respect, phasic, sound-evoked ACh release in the BLA that was selectively scaled up for the CS+ stimulus could facilitate the long-term synaptic plasticity from CAmy projections or more generally within the BLA that supports discriminative threat memory.

In previous studies, we noted a discriminative potentiation of spike rates and bulk calcium activity in cholinergic basal forebrain neurons that target the auditory cortex using trace conditioning and operant reinforcement learning protocols (Guo et al., 2019; Robert et al., 2021). Here, we found that ACh release was indiscriminately reduced after the first habituation day and was subsequently unchanged during threat memory acquisition and recall. The cholinergic basal forebrain neurons that target the primary auditory cortex are topographically distinct from the cholinergic neurons that target the higher-auditory cortex (Chavez and Zaborszky, 2016), which may speak to prior reports of categorically different learning-related plasticity within different regions of the cholinergic basal forebrain (Robert et al., 2021). Related to that point, our previous studies had targeted neural activity measured from the cholinergic cell bodies in the basal forebrain, whereas the fiber recordings described here measured ACh release at their postsynaptic targets. Therefore, the cholinergic neurons that contributed to the ACh release dynamics reported here are unknown and could be distinct from previous work that targeted particular regions of the basal forebrain. Further, ACh release and cholinergic neural activity based either on calcium imaging or spike recordings are not interchangeable. ACh binding to the fluorescent sensor competes with other endogenous ACh receptors and is itself shaped by acetylcholinesterase levels. Future experiments could disambiguate between these possibilities by performing the same type of neural activity measurement in the HO-AC and primary auditory cortex under various learning protocols where sound is associated with aversive reinforcement.

Although we did not perform ACh measurements in a separate cohort of pseudo-conditioned mice, several lines of reasoning argue against the necessity of this control experiment. Primarily, the fact that ACh release different in every way between the BLA and HO-AC (phasic, discriminative and generally potentiating in BLA, while more sluggish and non-discriminatively suppressed in HO-AC) is itself an internal control that argues against the involvement of an extraneous brain-wide contribution related to movement or another source of artifact. Secondly, the 405nm control wavelength captures small variations in signal that are unlikely related to ACh concentrations and this interleaved response was subtracted from the ACh3.0 sensor fluorescence signal. However, we cannot rule out the possibility that a globally reduction in signal to noise ratio could have contributed to the strong and non-specific habituation in sound-evoked ACh release in the HO-AC.

### Conclusion

Overall, we show that both corticoamygdalar as well as cholinergic inputs to BLA display discriminative forms of plasticity, mirroring the reorganization in CS encoding seen in BLA units and population responses. Future work using selective causal manipulations is needed to address whether these reorganized inputs are instructive signals guiding and maintaining the plasticity in BLA or whether they constitute a redundant encoding of memory distributed over distant brain regions. By regulating the persistence of plasticity in BLA, a maladaptive and overly persistent potentiation of corticoamygdalar or cholinergic inputs could also work against memory extinction, and hence teasing apart the role of these inputs could potentially provide additional insights into the neural underpinnings of PTSD and other anxiety related disorders.

## Acknowledgements

We thank Liam Casey for assistance with confocal microscopy, Ashwini Melkote for assistance with DeepLabCut, Ke Chen for sharing code on videography analyses, Christine Liu and Anne Takesian for advice on anatomical tracing and cell counting, and Sam Smith for guidance on data analysis. We thank Yulong Li for making the GRAB_ACh_3.0 sensor available for purchase. Financial support was provided by the Nancy Lurie Marks Family Foundation and NIH grants DC009836 and DC017078 (DBP).

## Author contributions

Conceptualization, M.M.A. and D.B.P.; Methodology, M.M.A., Y.W., E.Y.K. and D.B.P.; Investigation, M.M.A. and Y.W.; Software, M.M.A. and E.Y.K.; Formal Analysis, M.M.A.; Data Curation, M.M.A.; Visualization, M.M.A. and D.B.P.; Writing – Original Draft, M.M.A. and D.B.P.; Writing – Review & Editing, M.M.A. and D.B.P.; Resources, D.B.P.; Supervision, D.B.P.; Funding Acquisition, D.B.P.

## STAR ✶ Methods

### RESOURCE AVAILABILITY

#### Lead contact

Further information and requests should be directed to and will be fulfilled by the lead contact, Meenakshi Asokan.

#### Materials availability

This study did not generate new unique reagents.

#### Data and code availability

- All original data reported in this paper will be deposited at Mendeley Data and made publicly available as of the date of publication. DOIs will be listed in the key resources table.
- All original code will be deposited at GitHub and made publicly available as of the date of publication. DOIs will be listed in the key resources table.
- Any additional information required to reanalyze the data reported in this paper is available from the lead contact upon request.

### EXPERIMENTAL MODEL AND SUBJECT DETAILS

#### Animal subjects

We used adult male and female C57BL6 mice (Jackson Labs 000664) aged 9-10 weeks at the time of recording. Mice were housed individually after undergoing a major survival surgery. Mice were maintained in a 12/12 light/dark cycle with food and water available ad libitum and experiments were performed during their dark cycle. All procedures were approved by the Massachusetts Eye and Ear Infirmary Animal Care and Use Committee and followed the guidelines established by the National Institute of Health for the care and use of laboratory animals. Pupil- and facial motion-indexed behavioral measurements were performed in 27 mice, of which 3 were excluded for pupil dilation analysis because of pupil occlusion. Dual-site BLA and HO-AC electrophysiological recordings were performed in 11 of these mice; Dual-site BLA and HO-AC cholinergic sensor fiber recordings were performed in 8 of these mice, of which two were excluded from the analysis of BLA ACh levels because of imprecise placement of the fiber implant over BLA.

### METHOD DETAILS

#### Surgical preparation

Mice were anesthetized with isoflurane in oxygen (5% induction, 2% maintenance) and placed in a stereotaxic frame (Kopf Model 1900). A homeothermic blanket system was used to maintain body temperature at 36.6° (FHC). Lidocaine hydrochloride was administered subcutaneously to numb the scalp. At the conclusion of the procedure and 24hr post-recovery, Buprenex (0.05 mg/kg) and meloxicam (0.1 mg/kg) were administered, and the animal was transferred to a warmed recovery chamber.

#### Head plate attachment

The dorsal surface of the scalp was retracted and the periosteum was removed. The exposed skull surface was prepped with etchant (C&B metabond) and 70% ethanol before affixing a titanium head plate (iMaterialise) to the skull with dental cement (C&B Metabond). Mice were given at least 48 hours to recover, after which they were acclimated to the head fixation apparatus before the electrophysiological recordings.

#### Injections and fiber implantation

For all adeno associated viral vector (AAV) and retrograde tracer injections, mice were prepped as described above. For BLA injections, we first leveled the head by ensuring that the left and right z coordinates for the lateral skull were within +/− 0.03 mm and the z coordinate of lambda was within +/− 0.05 mm of bregma. For injections into the higher order auditory cortex (HO-ACtx), the temporalis muscle to expose the skull the over the right squamosal suture as it passes just dorsal to the rhinal fissure. Burr holes were made in the skull with a 31-gauage needle. Pulled glass micropipettes (Wiretrol II, Drummond) were backfilled with virus solution and injected into the target brain areas at 1 nl/s using a precision injection system (Nanoject III, Drummond) with a 5 or 8s delay between each injected bolus. BLA injection coordinates were 1.7 mm posterior from bregma (approximated intersection of skull sutures), 3.45 mm lateral of midline, 3.75 mm below the pial surface. HO-AC injection coordinates were 3.1 mm posterior to the bregma, lateral to the temporal ridge and medial to the squamosal suture, and 0.5 mm below the pial surface. At least 10 minutes passed following each injection before the pipette was withdrawn.

##### AAV injections to express ChR2 in HO-AC CAmy neurons

We injected 150 nl of AAVrg-pgk-cre (1.7 × 10^13^ genome copies/mL, Addgene 24593) into the BLA for retrograde expression of cre in the amygdala-projecting cells and 200 nl of AAV5-Ef1a-DIO-hChR2-EYFP (diluted 10% in sterile saline from 2.7 × 10^13^ genome copies/mL, Addgene 35509) into the HO-AC for cre-dependent hChR2 expression. We allowed 3 weeks for the virus incubation before performing electrophysiology experiments.

##### Retrograde tracing experiments

We injected 300 nl of Cholera Toxin Subunit B, Alexa Fluor 555 conjugate (CTB-AF555, Thermo Fisher Scientific) was injected into the BLA following the procedure above. Mice were euthanized 7-9 days after injections.

##### Express and measure the genetically encoded fluorescent ACh sensor

We injected 200 nl of AAV9-hSyn-ACh3.0 (diluted 2.5% in sterile saline from 3.6 × 10^13^ genome copies/mL, WZ Biosciences YL001003-AV9-PUB) into BLA and 300 nl of AAV9-hSyn-ACh3.0 (diluted 10% in sterile saline from 3.6 × 10^13^ genome copies/mL, WZ Biosciences YL001003-AV9-PUB) into HO-AC. An optic fiber was implanted following each injection such that the distal tip of the fiber terminated 0.15 – 0.25mm above the injection depth. We implanted a flat fiber into BLA (Doric, NA 0.37, 2mm length) and an angled fiber (Doric, NA 0.37, 5mm length, 45 deg angle) into HO-AC. Both fibers featured a 0.2mm core diameter and zirconia ferrule receptive (outer diameter 1.25mm). Fibers were fixed into place and optically sealed by applying dental cement mixed with black India Ink to the exposed skull and head plate. We allowed 3 weeks for the virus incubation before performing bulk fiber measurements.

#### Electrophysiology

##### Preparation for acute insertion of high-density probes in awake, head-fixed mice

A ground wire was implanted atop the left occipital cortex via a small burr hole during the preceding head plate attachment procedure described above. On the day of the Habituation session, mice were briefly anesthetized with isoflurane in oxygen (5% induction, 2% maintenance) and two small (~1 × 1-1.5 mm) craniotomies were made in the right hemisphere using a scalpel, each centered on the prior injection location. A circular well was constructed around each craniotomy with UV-cured cement (Flow-It ALC Flowable Composite) and filled with lubricating ointment (Paralube Vet Ointment) and the isoflurane was discontinued. Mice were placed in a body cradle and their head was immobilized by attaching the headplate to a head fixation post. Recordings were performed inside a dimly lit single-wall sound attenuating recording chamber (Acoustic Systems) after allowing at least 30 minutes to fully recover from anesthesia. At the conclusion of each recording session, the craniotomy was flushed with saline, ointment re-applied, and the recording well was sealed with a cap of UV-cured cement.

##### Extracellular recordings

BLA recordings were performed with a two-shank 64-channel silicon probe (Cambridge Neurotech; H2 probe, 25 μm spacing between contacts within a shank, and 200μm spacing between shanks). HO-AC recordings were made with a single shank optrode (Cambridge Neurotech; H3 probe with 20μm spacing between contacts. The attached optic fiber featured a flat tip (200μm core, 0.66 NA), a 200μm horizontal offset to the shank, and 125μm vertical offset between the fiber tip and most superficial channel. Probes were positioned with a micromanipulator (Narishige) and inserted via a hydraulic microdrive (FHC). HO-AC recordings were made with an oblique insertion angle that spanned AuV and TeA. The BLA recording probe was lowered ventrally with the two shanks oriented medio-laterally while optogenetically activating CAmy neurons (473nm diode laser, Omicron, LuxX) with brief laser pulses (1 ms duration, 10 Hz, 20 mW) to identify light-activated multiunit activity. The distal tip of the BLA recording probe was typically 3.7-4mm below the pial surface but the fine position was adjusted such that CAmy-evoked multiunit responses were nearly absent in the most ventral channel. Once both probes were in place, the brain settled for approximately 15 mins before recordings began.

#### Fiber photometry

LEDs of different wavelengths provided a basis for separating ACh-dependent fluorescence (465nm) from ACh-independent (405nm) fluorescence. LEDs were modulated at 210Hz (465nm) and 330Hz (405nm), respectively, and combined through an integrated fluorescence mini-cube (FMC4, Doric). The optical patch cable was connected to the fiber implant via a zirconia mating sleeve to produce a tip power of 0.1 - 0.2mW. Bulk fluorescent signals were acquired with a femtowatt photoreceiver (2151, Newport) and digital signal processor (Tucker-Davis Technologies RZ5D). The signal was demodulated by the lock-in amplifier implemented in the processor, sampled at 1017Hz and low-pass filtered with a corner frequency at 20Hz. The optical fibers were prebleached overnight by setting both LEDs to constant illumination at a low power (<50μW).

#### Discriminative threat conditioning under head-fixation

DTC was performed in three phases: Habituation, Conditioning and Recall. For electrophysiology recordings, each phase was performed in a single daily session separated by approximately 24 hours. For fiber recordings, the Habituation phase was two sessions, Conditioning was three sessions, and Recall one session, each separated by approximately 24 hours. Parameters for DTC including CS and aversive reinforcement were based on recent publications (Belén Pardi et al., 2020; Dalmay et al., 2019). All sessions presented trains of five frequency modulated (FM) sweeps presented at 1Hz (0.5s duration, 70 dB sound pressure level, 50ms raised cosine onset and offset gating applied at the FM endpoints). Each of the five FM sweeps for a given trial either increased or decreased in frequency (5-20kHz or 20-5kHz, respectively) at a rate of 4 octaves/sec. Daily sessions consisted of 30 alternating upward FM or downward FM trials with a 20-180s inter-trial interval selected from a decaying exponential distribution to produce a flat hazard function. On Conditioning sessions, one FM sweep direction, the CS+, the 5^th^ FM sweep coincided with the onset of a mildly aversive tail-shock (1 s, 0.4 mA AC, Coulbourn Precision Animal Shocker) via pediatric cuff electrodes positioned ~1 cm apart at the center of the tail. The assignment of the CS+ FM sweep direction was counterbalanced between animals. The cradle and surrounding test apparatus was cleaned with 70% ethanol before Habituation and Conditioning sessions, and 0.2% acetic acid before the Recall session.

Pseudo-conditioning was performed identically, except that the timing of the 15 tail shocks, 15 upward FM sweep trains, and 15 downward FM sweep train were each separated by the 20-180 s inter trial interval. Audio stimuli were generated with a 24-bit digital-to-analog converter (National Instruments model PXI-4461), and presented via a free-field speaker (Parts Express 275-010) placed approximately 10 cm from the left (contralateral) ear canal. Free-field stimuli were calibrated using a wide-band free-field microphone (PCB Electronics, 378C01).

#### Pupillometry and facial videography

Video recordings of the pupil and face were acquired at 30Hz with a CMOS camera (Teledyne Dalsa, model M2020) outfitted with a lens (Tamron 032938) and infrared longpass filter (Midopt lp830, 25.5nm cutoff). Recordings were made in isoluminous lighting provided by infrared LEDs (850 nm, Vishay Semiconductors, VSLY5850) where additional ambient light in the visible spectrum was adjusted to maintain an intermediate steady state pupil diameter.

#### Histology

Deeply anesthetized mice were perfused transcardially with 0.01 M phosphate-buffered saline (PBS; pH = 7.4) followed by 4% formaldehyde in 0.01 M PBS. Brains were removed and stored in 4% formaldehyde for 12 h before transferring to cryoprotectant (30% sucrose in 0.01 M PBS) for at least 48 hrs. Coronal sections were cut at 40 μm thickness on a cryostat and coverslipped using Vectashield Mounting Medium with DAPI (Vector Labs). Sections were imaged with a 10X/0.40 NA dry objective using an epifluorescence microscope (Leica DM5500B) or under a 40X /1.30 NA oil immersion objective using a confocal laser scanning microscope (Leica SP8).

### QUANTIFICATIONS AND STATISTICAL ANALYSIS

#### Electrophysiology data acquisition and online analysis

Raw neural signals were digitized at 32-bit, 24.4 kHz and stored in binary format (PZ5 Neurodigitizer, RZ2 BioAmp Processor, RS4 Data Streamer; Tucker-Davis Technologies). To eliminate artifacts, electrical signals were notch filtered at 60 Hz, the Common-mode signal (channel-averaged trace) was subtracted from the raw signals from all channels, independently for each probe. For online visualization, signals were band-pass filtered (300-3000 Hz, second-order Butterworth filters) and multiunit activity was extracted as negative deflections in the electrical trace with an amplitude exceeding 4 standard deviations of the baseline hash.

#### Single unit identification and analysis

##### Single unit isolation

We used Kilosort2 (Pachitariu et al., 2016) to sort spikes into single unit clusters. For recordings done on Habituation and Recall sessions, we concatenated all data files from a given session so that the same unit could be tracked over the full course of the experiment (~90 min). We ensured our units were isolated clusters with inter-spike intervals > 2ms for at least 95% of all spikes. Once isolated, spike waveforms with trough to peak intervals > 0.6ms were as regular spiking putative excitatory neurons, while intervals < 0.6ms were classified as fast spiking putative inhibitory interneurons, as per our previous work (Asokan et al., 2021).

##### Optogenetic identification of corticoamygdalar units

We operationally defined units with a high laser evoked spiking rate (> 5 standard deviations above prestimulus baseline), low first spike latency (< 5 ms) and a low first spike jitter (standard deviation of first spike latency < 0.75ms) in response to a 1ms 20mW laser pulse stimulation presented at 1Hz as the photo-identified corticoamygdalar cells.

##### Analysis of evoked firing rate and stimulus synchrony

Only neurons that fired at least 0.01 Hz across the whole session were included for analysis. CS− evoked firing rates were measured in units with a peak firing rate >1.5 standard deviation above pre-stimulus baseline during the post-stimulus response period for the first FM sweep of the train for either CS, as determined with 1ms binning. The CS− evoked response used for computing the discriminative plasticity was quantified as the area under the curve of the peri-stimulus time histograms (PSTH) over the 5s CS duration. Asymmetry indices were computed as 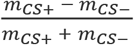 where *m* is a measure with positive values such as neural firing rate (in spikes/s). Stimulus synchrony of the units was visualized using the firing rate averaged over 30 upward and downward sweeps and expressed as a z-score with respect to the baseline firing during a 1s duration before the sound onset. It was quantified as the amplitude at 1Hz of the fast Fourier transform of this z-scored firing rate during the 5s CS duration.

#### Dimensionality reduction and neural population trajectories

Trial-averaged spike rates were expressed as z-scores relative to the distribution of pre-stimulus firing rates and smoothed with a 100ms gaussian filter. We then constructed a matrix with the concatenated responses to each CS for each unit on a row. The mean response for each unit was then subtracted from all column values. We performed singular value decomposition on this matrix using the Matlab function ‘svd’, and obtain its projections onto the transformed subspace, thereby reducing dimensionality using principal component analysis. To visualize the neural population trajectories, we plot the temporal evolution of the responses to each CS in the space defined by the first three principal components. To compute the Euclidian distance between the CS trajectories, we use the number of dimensions that are necessary to explain 80% of the variance in the data.

#### Evoked LFP amplitudes and Spike-triggered LFPs

##### LFP extraction

To extract the LFPs, raw signals from each channel of the recording electrodes were notch filtered at 60 Hz, down-sampled to 1000 Hz and spatially smoothed with a triangle filter (5-point Hanning window). We then subtracted the Common-mode reference (average signal across all channels) from each channel.

##### Analysis of evoked LFP amplitudes

CAmy-evoked LFP response in BLA was measured by stimulating the CAmy cells using 100 repetitions of a 1ms 0-20 mW laser pulse presented at 4Hz. The CAmy-evoked LFP amplitude for each channel in BLA was computed as the absolute value of peak of the deflection of the averaged LFP response in the 50ms duration following the laser pulse. The sound-evoked LFP amplitudes in each channel were expressed as the average instantaneous amplitude during the 5s CS duration calculated from the amplitude of the complex Hilbert transform of the LFP.

##### Spike-triggered LFPs

Network-level functional coupling was estimated from the spike-triggered LFP (stLFP). To estimate the stLFP, we used linear deconvolution by time expansion. The LFP measured in one region, for example BLA (y) was interpreted as a sum of the linear convolution of the spiking events in the other region, for example HO-AC with the isolated (HO-AC → BLA) stLFP (ß), and all the other possible sources (e) in:

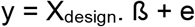

Deconvolution was used to recover the unknown stLFP given only the measured LFP and the time of the spiking events (which is used to construct the design matrix), and estimate the stLFP that best explain the observed LFP given the spike times (Ehinger and Dimigen, 2019). The spiking events can occur at any temporal interval between each other, and it is assumed that their contributions to y will linearly add up. We created a time-expanded version of the design matrix (X_design_) with several time points around each event added as predictors and we then solved the model for the stLFP. The stLFP evoked by each unit was averaged across all channels in the other region. Since the sign of the stLFP deflection varies across depth along the probe after common-mode referencing, positive-deflecting traces were inverted before averaging (Laszlovszky et al., 2020).

#### Photometry signal pre-processing and analyses

We calculated the ACh3.0 responses as the percentage fractional change in fluorescence ΔF/F_0_ (%), where F_0_ was defined as the running median fluorescence value in a 60 s time window. To reduce the potential contribution of intrinsic signals and movement artifacts, analyses were performed on a corrected ACh3.0 signal in which the fractional change in fluorescence measured with the 405nm excitation was smoothed using a 1 s gaussian filter and then subtracted from the 465nm signal for each trial (Rajebhosale et al., 2021). The CS− evoked responses were then quantified as the area under the curve of these corrected ACh3.0 signal during the initial 4s CS period.

#### Pupil dilation response

Pupil diameter (P) was measured with DeepLabCut (version 2.0) (Mathis et al., 2018; Nath et al., 2019). We labeled the four cardinal and four intercardinal points for the right pupil of each mouse in 10 frames each from 31 animals recorded under similar conditions. Training was performed on 95% of frames. We used a ResNet-101 based neural network with default parameters for 1.03 million training iterations. Each tracked point was expressed as a 3-D vector as × coordinate × y coordinate × time. Pupil diameter was estimated from the distance between East – West markers, which proved most robust to variations in eye lid position. Frames with the likelihood of these markers < 0.7 were discarded (e.g., during blink) and values were determined by interpolation. The sound-evoked pupil dilation was computed as a fractional change in pupil diameter (ΔP/P_0_) with respect to the mean pupil diameter at baseline (P_0_, 2s before sound onset).

#### Facial motion response

As per previous work (Stringer et al., 2019), facial motion energy was measured at time T as the absolute value of the difference in pixel intensities between consecutive frames (T, T+1) for each pixel within the region of interest. We then positioned a region of interest (ROI) on the rostral cheek, just caudal to the vibrissae array, and defined facial motion (F) as a sum of the total motion energy for all pixels within the region of interest. We then expressed the sound-evoked facial motion for each trial as a fractional change (ΔF/F_0_) with respect to the mean facial motion in the baseline (F_0_, 2s before sound onset).

#### Cell count quantification and electrode tracks reconstruction from photomicrographs

To count CTB-labeled cells in the auditory cortex, we first used SHARP-Track (Shamash et al., 2018) to register the photomicrographs with the Allen brain atlas. The center of each labeled cell was then marked with image processing software (Fiji) and the coordinates of each point was saved. We then marked the pial surface as a line and ran a function (developed by Michael Cammer, Microscopy core NYU Langone Medical Center) to the shortest distance of all the cells from the indicated pial surface.

To validate the electrophysiology recording locations, probe shanks were dipped in a fluorescent lipophilic dye (DiI, Sigma-Aldrich 42364) before the final recording session and their insertion paths reconstructed from post-mortem photomicrographs using SHARP-Track (Shamash et al., 2018).

#### Statistical analysis

All statistical analysis was performed with MATLAB (Mathworks). Non-parametric statistical tests were used in cases where data samples did not meet the assumptions of parametric statistical tests. Effect sizes were estimated with Cohen’s d for normally distributed data and with Cliff’s delta for samples that did not conform to a normal distribution. We used the standard p-value < 0.05 for assigning statistical significance denoted by asterisk symbol. The standard p-value was used in conjunction with a Cohen’s d > 0.4 (or Cliff’s delta > 0.3, which are traditionally assigned to a medium-sized effects or greater) in cases where the sample size was high (>25). Multiple post-hoc comparisons were corrected using Bonferroni-Holm correction.

## Supplemental information

**Supplemental Figure 1:**
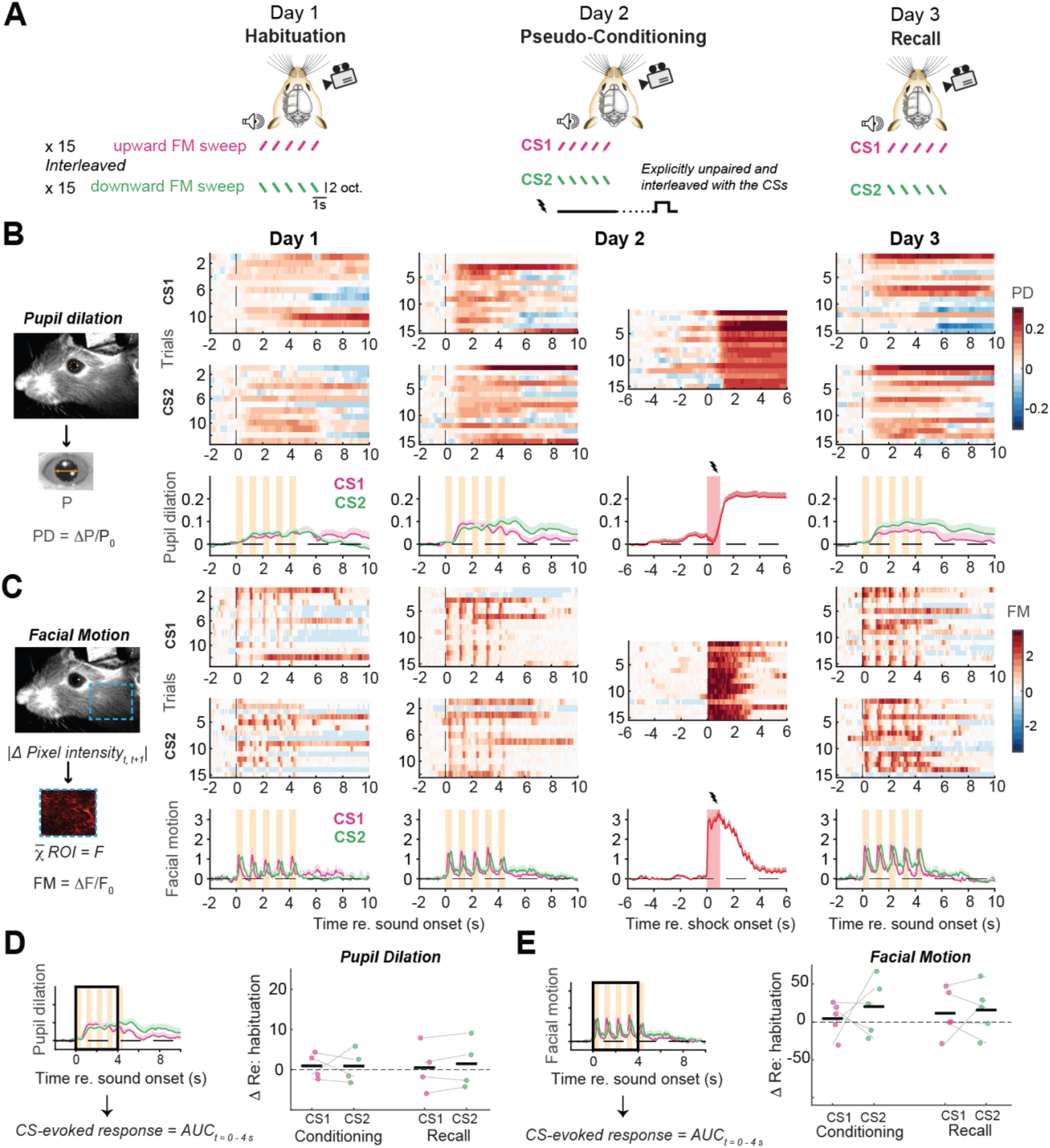
Pseudo-conditioned mice do not show any discriminative or generalized changes in pupil dilation and facial motion. **A**) Schematic illustrating the Pseudo-conditioning protocol, where each of the three sessions are separated by 24 hours. In all the sessions, the mice are presented with 15 alternating presentations of a train of frequency modulated (FM) sweep in upwards or downwards direction (CS1 and CS2) during high-resolution facial videography. In the conditioning sessions, a mildly aversive tail-shock is randomly interleaved and presented in an explicitly unpaired fashion. **B**) *Left:* Pupil dilation in each trial is quantified as a fractional change in the pupil diameter (P) with respect to the mean pupil diameter in the 2s baseline before CS onset (ΔP/P_0_). *Right:* Pupil dilation for all 15 presentations of CS1, CS2, tail-shock (*top and middle*), and mean pupil dilation across trials (*bottom*) for all three sessions in an example mouse. Vertical dashed lines denote onset of initial FM sweep, orange bars denote CS duration, and red bars denote the 1s shock. **C**) *Left:* Facial motion is computed at each time T as the absolute value of the difference in pixel intensities between consecutive frames (T, T+1) for each pixel and averaged over all the pixels within the region of interest (dashed blue rectangle). *Right:* Facial motion was expressed as a fractional change with respect to the mean facial motion in the 2s baseline before CS onset (ΔF/F_0_). Other plotting conventions match above. **D**) *Left:* Illustration of the area under the curve (AUC) quantification approach for pupil diameter during the first 4 seconds of the CS presentation (black rectangle). *Right:* Difference in mean CS+ and CS− AUC for pupil dilation during Conditioning and Recall sessions relative to Habituation. Horizontal black bars indicate the mean. Pupil dilations do not systematically differ by CS type or Session (N = 4 mice): 2-way repeated measures ANOVA, main effect for Session [F = 1.000, p = 0.500], Sound [F = 1.000, p = 0.500], Session × Sound interaction [F = 0.269, p = 0.632]. **E**) Same plotting conventions as D but for Facial motion. Facial motion also does not systematically differ by CS type or Session (N = 5 mice): 2-way repeated measures ANOVA, main effect for Session [F = 1.000, p = 0.500], Sound [F = 1.000, p = 0.500], Session × Sound interaction [F = 0.214, p = 0.675].

**Supplemental Figure 2:**
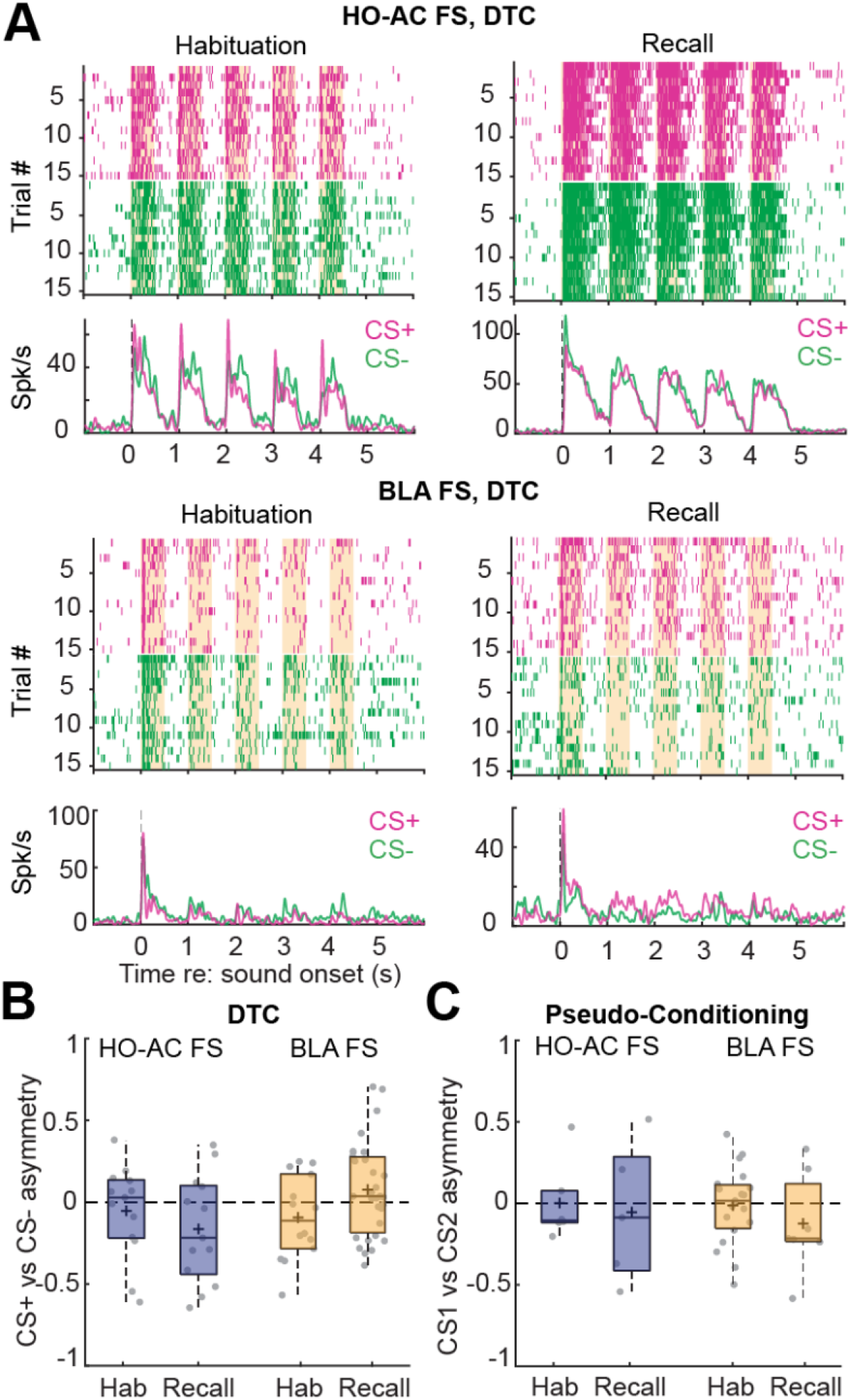
Fast-spiking units from HO-AC and BLA do not show a significant bias in the CS− evoked responses with DTC or Pseudo-conditioning. **A**) Rastergrams and peri-stimulus time histograms from example HO-AC and BLA FS units recorded on the Habituation and Recall sessions of DTC. **B**) Discriminative plasticity from sound-responsive units in 8 mice that underwent DTC using an asymmetry index ((CS+ – CS−) / (CS+ + CS−), where positive values reflect a greater response to the CS+, negative values to the CS− and a value of zero reflects an equivalent response to both stimuli. CS− evoked responses were not biased towards the CS+ in the Recall session compared to Habituation in HO-AC FS units (n = 13/13 Habituation/Recall; unpaired t-test, p = 0.37, Cohen’s d = −0.36) and were only marginally biased towards the CS+ in BLA FS units (n = 14/31 Habituation/Recall; p = 0.07, Cohen’s d = 0.61). **C**) Discriminative plasticity from sound-responsive units in 3 mice that underwent Pseudo-Conditioning with the same analysis described above. CS− evoked responses did not show a significant difference in bias in HO-AC FS units (n = 6/5 Habituation/Recall; unpaired t-test, p = 0.79, Cohen’s d = −0.17) or BLA FS units (n = 19/7 Habituation/Recall; p = 0.34, Cohen’s d = −0.43).

**Supplemental Figure 3:**
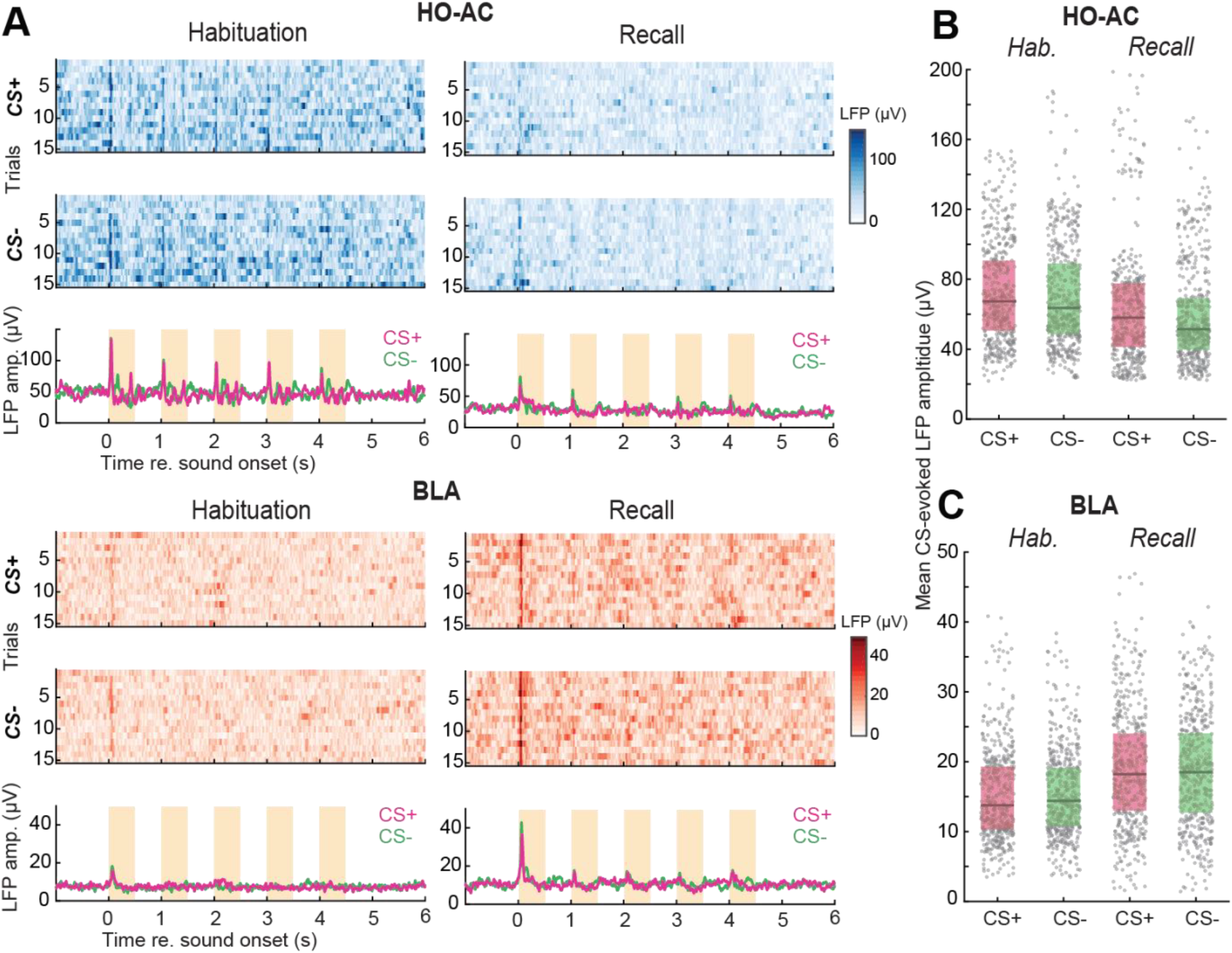
Sound-evoked LFPs in HO-AC and BLA do not show a CS+ response bias. **A**) The instantaneous amplitude of the LFP in example channels of HO-AC and BLA for all 15 presentations of CS+ and CS− (*top and middle*) and the mean LFP amplitude across the trials (*bottom*) during Habituation and Recall in a mouse that underwent DTC. Orange bars denote CS duration. **B**) Mean CS− evoked instantaneous amplitude of the HO-AC LFP during the 5s CS duration for all recording channels during Habituation and Recall in mice that underwent DTC (N = 8/512 mice/channels). HO-AC LFP amplitudes are suppressed during Recall (Mixed model ANOVA with Session as a factor and Sound as a repeated measure, main effect for Session [F = 10.46, p = 0.0013], main effect for Sound [F = 75.96, p = 1.15 x10^−17^], Session × Sound interaction [F = 36.34, P = 2.31 × 10^−9^]), and the CS− specific differences are marginal with the effect size (Cohen’s d) taken into account (paired t-tests, p < 0.008 but Cohen’s d < 0.4 for Habituation and Recall). **C**) Same as above, but for all BLA channels (N = 8/512 mice/channels). BLA LFP amplitudes are enhanced during Recall (Mixed model ANOVA with Session as a factor and Sound as a repeated measure, main effect for Session [F = 37.051, p = 1.63 × 10^−9^], main effect for Sound [F = 0.046, p = 0.830], Session × Sound interaction term [F = 9.223, P = 0.0025]), but there is no response bias towards CS+ after DTC with the effect size taken into account (paired t-tests, Habituation, p value/Cohen’s d = 0.01/0.13; Recall, p- alue/Cohen’s d = 0.09/-0.08).

**Supplemental Figure 4:**
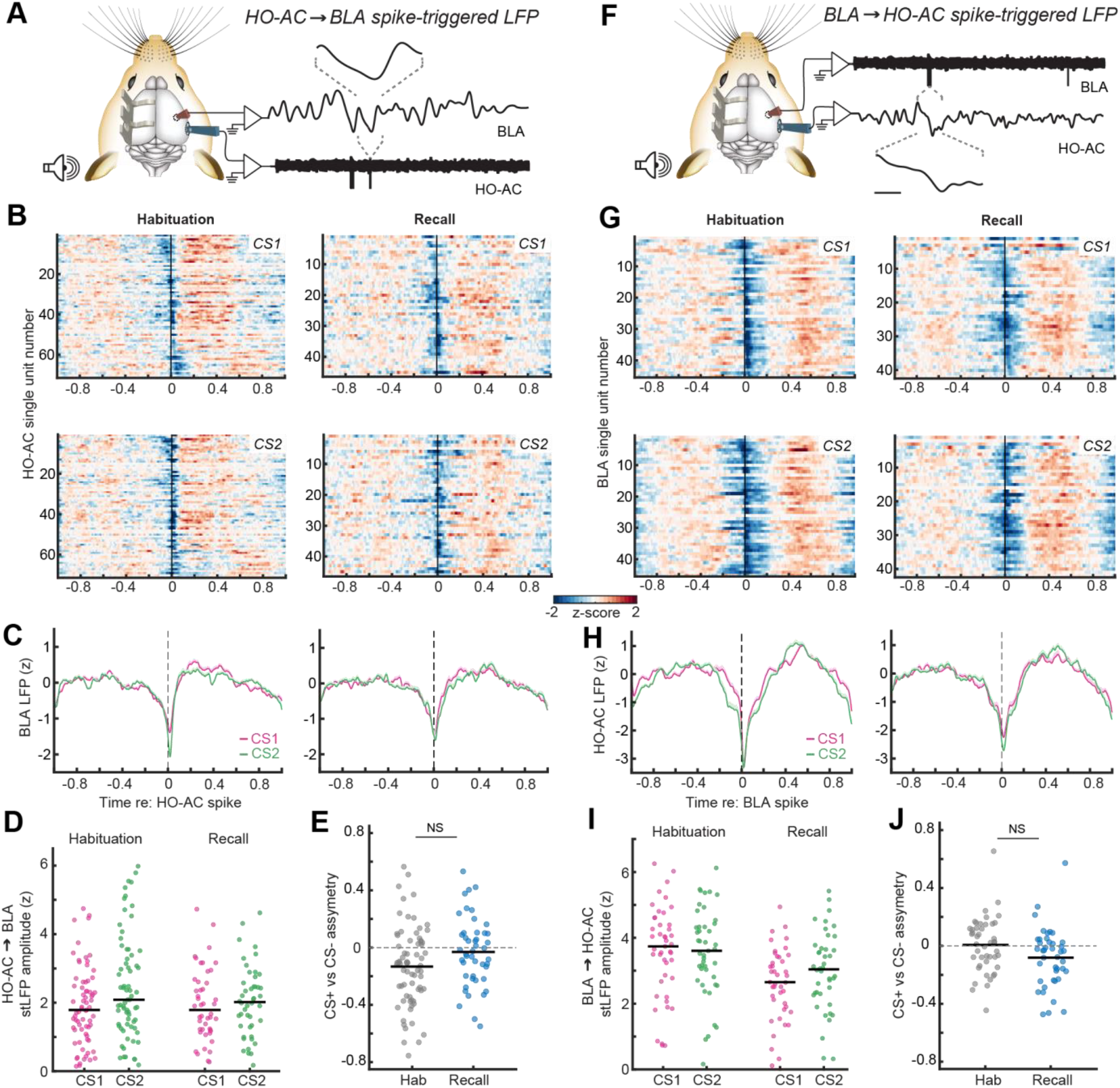
Pseudo-conditioned mice do not show changes in functional coupling from HO-AC to BLA or BLA to HO-AC. **A**) Schematic illustrating the quantification of BLA LFPs triggered by HO-AC single unit spikes. Linear deconvolution by time expansion is used to estimate the spike-triggered LFP (stLFP). **B**) Estimated HO-AC to BLA stLFPs computed during the CS1 (*top*) and CS2 (*bottom*) expressed as a z-score relative to pre-stimulus baseline and averaged across all recording channels in BLA in mice that underwent Pseudo-conditioning. **C**) Mean ± SEM HO-AC to BLA stLFP demonstrates a downward deflection of the BLA shortly following HO-AC spikes for both the CSs but no CS− specific difference after Pseudo-conditioning. **D**) BLA stLFP amplitude for each HO-AC RS unit during each CS presentation on Habituation and Recall sessions (N/n = 3/71, 3/46 mice/units for Habituation and Recall respectively). Horizontal black bars indicate the mean. BLA stLFP is elevated during CS2 stimuli during Habituation but the CS− specific difference is absent during Recall: Mixed model ANOVA with Session as a factor and Sound as a repeated measure, main effect for Session [F = 1.131, p = 0.290], main effect for Sound [F = 11.005, p = 0.001], Session × Sound interaction term [F = 5.652, P = 0.019]. **E**) Discriminative plasticity in the HO-AC to BLA stLFP for each unit can be expressed as an asymmetry index ([CS+ - CS−] / [CS+ + CS−] where positive values reflect a greater response to the CS+, negative values to the CS− and a value of zero denotes an equivalent response. The asymmetry index was on average negative during Habituation (one-sample t-test, p value/Cohen’s d = = 0.0004/0.48), but was neither significantly biased in the Recall session (one-sample t-test, p value/Cohen’s d = 0.39/-0.13) nor significantly different than in Recall compared to Habituation (unpaired t-test, p value/Cohen’s d = 0.09/0.39). Horizontal black bars indicate the mean. **F-H**) As per *A-C*, but for the HO-AC LFP triggered by spikes in individual BLA units. **I**) Plotting conventions match *D*. HO-AC stLFP amplitude for each BLA unit (N/n = 3/45, 3/42 mice/units for Habituation and Recall respectively). Horizontal black bars indicate the mean. HO-AC stLFP was reduced during Recall with no significant CS− specific changes: Mixed model ANOVA with Session as a factor and Sound as a repeated measure, main effect for Session [F = 9.732, p = 0.002], main effect for Sound [F = 2.596, p = 0.111], Session × Sound interaction term [F = 3.208, p = 0.077]. **J**) Plotting conventions match *E*. BLA to HO-AC stLFP amplitude was not significantly biased with the effect size taken into account (one-sample t-test, Habituation, p value/Cohen’s d = 0.835/0.031; Recall, p value/Cohen’s d = 0.042/-0.397), and was not significantly different than in Recall compared to Habituation (Unpaired t-test, p value/Cohen’s d = 0.087/-0.439). Horizontal black bars indicate the mean.

